# Simultaneous integration of gene expression and nutrient availability for studying metabolism of hepatocellular carcinoma

**DOI:** 10.1101/674150

**Authors:** Ewelina Weglarz-Tomczak, Thierry D.G.A. Mondeel, Diewertje G.E. Piebes, Hans V. Westerhoff

## Abstract

How cancer cells utilize nutrients to support their growth and proliferation in complex nutritional systems is still an open question. However, it is certainly determined by both genetics and an environmental-specific context. The interactions between them lead to profound metabolic specialization, such as consuming glucose and glutamine and producing lactate at prodigious rates. To investigate whether and how glucose and glutamine availability impact metabolic specialization, we integrated computational modeling on the genome-scale metabolic reconstruction with an experimental study on cell lines. We used the most comprehensive human metabolic network model to date, Recon3D, to build cell line-specific models. RNA-Seq data was used to specify the activity of genes in each cell line and the uptake rates were quantitatively constrained according to nutrient availability. To integrated both constraints we applied a novel method, named GENSI (Gene Expression and Nutrients Simultaneous Integration), that translates the relative importance of gene expression and nutrient availability data into the metabolic fluxes based on an observed experimental feature(s). We applied GENSI to study hepatocellular carcinoma addiction to glucose/glutamine. We were able to identify that proliferation, and lactate production is associated with the presence of glucose but does not necessarily increase with its concentration when the latter exceeds the physiological concentration. There was no such association with glutamine. We show that the integration of gene expression and nutrient availability data into genome-wide models improves the prediction of metabolic phenotypes.

## 1. Introduction

Cancer cells adapt their metabolism to promote fast cellular proliferation and long-term maintenance [1–4], thus facilitating the uptake and conversion of nutrients into biomass. Most metabolic signatures are shared across different kinds of cancer cells, including one of the best recognizable, namely changes in glucose metabolism that give rise to the Warburg effect [1–7], and an increase in biosynthetic activities (such as nucleotides, lipids, and amino-acids synthesis). However, the Warburg effect also plays an important role in many other cell types involved in immunity, angiogenesis, pluripotency, and infection by pathogens [8]. The Warburg effect is characterized by an increased rate of glucose uptake and part of glycolysis-derived pyruvate is diverted to lactate [1–9], which produces much less ATP per glucose than its oxidation to carbon dioxide would. This metabolic signature enables fast-dividing cells to satisfy anabolic needs for biomass production and is accompanied by a suppression of apoptotic signaling [10–13]. The high glucose consumption during the Warburg effect also provides a higher production of reduced nicotinamide adenine dinucleotide phosphate (NADPH2) through the pentose phosphate pathway, which provides electrons for cell proliferation [14]. However, it does mean that oxidative metabolism (i.e., respiration) is damaged. Instead, respiration and other mitochondrial activities are required for tumor growth [15,16]. Many cancer cells also have an increased uptake of glutamine [17–21]. The partial catabolism of this glutamine to lactate by cancer cells has been called the WarburQ effect [22]. Some rapidly proliferating cells are particularly dependent on glutamine, and undergo necrosis upon glutamine depletion [20].

Genome-scale metabolic models (GEMs), which are at the core of some bottom-up systems biology approaches, can be used to predict cell physiology (e.g., growth rate and metabolic fluxes) under different conditions and improve our understanding of cell metabolism [23,24]. Since GEMs utilize a Boolean formulation connecting genes to reactions, they have been used extensively as platforms for analyzing mRNA expression data to elucidate how changes in gene expression impact cellular phenotypes [25–37]. There are two fundamental approaches for integrating gene-expression data into GEMs: (1) based on direct integration of the gene expression information into the flux bounds and (2) based on a categorization of genes. The first way includes, for example, setting the fluxes to zero if an expression of their associated genes was low [27] and the maximum allowable flux value as a function of measured gene expression [28]. In the second approach the reactions are divided into different categories based on gene expression (e.g. highly or lowly expressed) and then reactions with highly expressed genes are associated with high flux and reactions with lowly expressed genes with non-high flux. Such an approach had been applied in Gene Inactivity Moderated by Metabolism and Expression (GIMME) tool [29] where minimizes the flux through reactions whose associated genes’ expression falls below a given threshold. In contrast, Shlomi et al., in their Integrative Metabolic Analysis Tool (IMAT) divided the reactions into those associated with highly expressed genes and those associated with lowly expressed genes and then maximized the number of reactions whose fluxes are consistent with their gene expression state [30]. Graudenzia et al. introduced the data integration framework named Metabolic Reaction Enrichment Analysis (MaREA) by projecting RNA-seq data onto metabolic networks by assigning a score for each reaction in the network (Reaction Activity Score, RAS). The score is calculated based on the expression of genes encoding for the associated enzyme(s) [31]. Such a methodology is highly useful due to the fact that metabolic reconstruction of higher organisms does not have a one-to-one relationship between genes and network edges, due to the existence of isozymes and protein complexes.

However, genetics is not the only determinant of metabolic phenotype. Cancer cell metabolism, similar to any other type of cell, is also influenced by metabolic constraints imposed by environmental and tissue-specific contexts [16,39–49]. Numerous microenvironmental factors influence cancer cell metabolism [39–41,45,49] including tumor acidity [51,52] that is directly connected with lactate secretion and tumor nutrient levels [42,43,48]. In particular, environmental nutrient availability (NA) is an important regulator of cancer cell metabolism, and therefore an important environmental determinant of cancer cell metabolism is diet, which can affect the availability of nutrients within tumours [39,40,48,50].

However, including concentrations of nutrients into GEMs in order to study their impact on cancer cell metabolism is not yet widely implemented. The existing methods for including nutrients concentrations into GEMs, which are based on the correlation of exchange fluxes with relative estimates of consumption and/or secretion rates, were applied to study the metabolic adaptation of Escherichia coli in complex nutritional systems [53,54].

In the present study, we simultaneously constrain GEM and integrate linearly both: cell-intrinsic factors (gene expression level) and cell-extrinsic factors (Nutrient Availability). We applied a novel method, named GENSI (Gene Expression and Nutrients Simultaneous Integration), that translates genes expression and nutrient availability data into the fluxes through a scaling factor. The maximum allowable flux value of reactions associated with genes was constrained as a function of RAS [31] calculated based on gene expression level. While the maximum allowable flux value of the exchange reaction was constrained as a function of the possible consumption rate of the available nutrients. We integrated those constraints through Flux Balance Analysis [55] and a ‘scaling factor’. Using GENSI we prepared specific models of the two hepatocellular carcinoma cell lines, HuH7 and PLC/PRF/5, and we used these to predict the influence of glucose and glutamine availability on proliferation rate and some cancer-related metabolic behaviors. We were able to identify glucose and glutamine (in)dependencies of both cell lines. Predictions were then confirmed experimentally.

## 2. Results

### 2.1. GENSI methodology

We designed GENSI as a method to integrate relative gene-expression and nutrient availability data into the human genome-wide metabolic reconstruction. Our purpose was reducing the solution space of optimal fluxes to provide results that can predict cell physiologies based on both cell-intrinsic and cell-extrinsic factors. Cancer cell metabolism, which aroused our interest, is also influenced by metabolic constraints imposed by environmental contexts. One of the important environmental determinants of cancer cell metabolism is diet, which can affect the availability of nutrients within tumours.

The GENSI workflow along with an illustration of the method performed on a GEM model is presented in Figure 1. GENSI requires three inputs: 1) a GEM model, 2) gene-expression data and 3) nutrient availability data (Figure 1).

**Figure 1.**
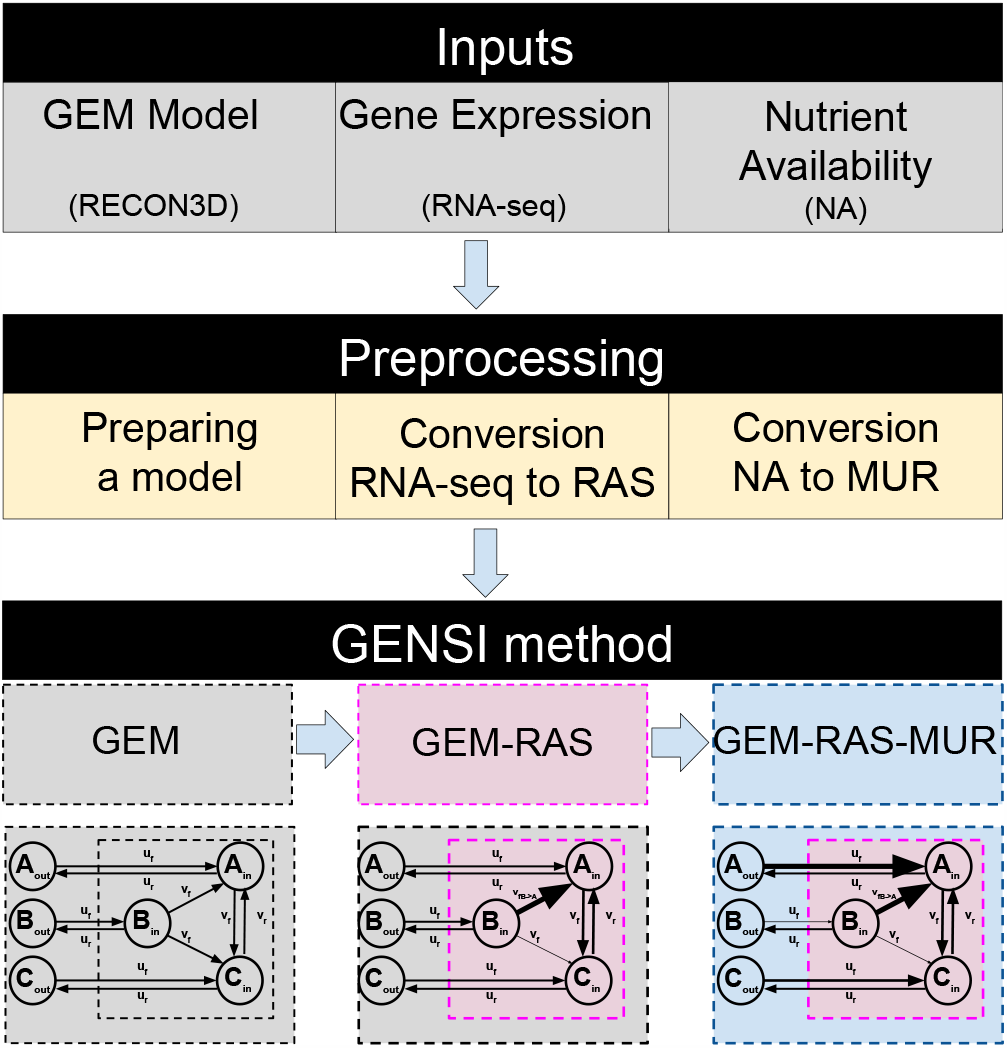
The GENSI workflow requires three inputs: a GEM model, gene-expression, and nutrient availability data (first block). In the pre-processing step (second block), the RNA-seq is converted into RAS [31] that is defined for any reaction r associated with gene(s) in the GEM and describes the extent of its activity, as a function of the expression of the genes encoding for the subunits and/or the isoforms of the associated enzyme(s). The NA data is converted to a Maximal Uptake Rate (MUR) that is defined for any exchange reaction *rex* in the GEM and describes the rate of the maximum possible uptake over time for substances available for the model (nutrients). Substances available for uptake are limited by the composition of the medium. Finally, GENSI (third block) integrates GEM, RAS, and MUR.

The first step of the GENSI framework consists of pre-processing of the GEM model and includes: 1) preparing the model and includes blocking of the uptake fluxes of various metabolites that are not present in NA, conversion of the gene identifiers that they are compatible with symbols used in RNA-seq data; 2) conversion of the gene-expression levels into RAS score [31]; and 3) conversion of the NA data into Maximal Uptake Rate (MUR) (Figure 1).

In the next step GENSI simultaneously translates RAS and MUR into the fluxes using a scaling factor of the RAS that is adjusted until experimentally observed features appear. As the direct correlation of gene expression data with maximal flux is not possible to apply in human metabolic map RECON3D [38], due to the existence of isozymes and protein complexes, we assigned to each reaction a RAS by summing over isoenzymes (for OR logic) and taking minima of subunits of a complex (for AND logic) of the TPM scores for the genes coupled to each reaction from the pre-processing step [28,31,56]. In this way, isoenzymes are thought to contribute additively to the activity of a reaction whereas the lack of even one subunit of an enzyme complex can bring down a reaction’s activity [31]. To integrate nutrient availability data into the GEM model we propose using Maximal Uptake Rate (MUR). We defined MUR for all exchange reactions in the GEM. It describes the rate of maximum possible uptake over time for substances (nutrients) available for the cells.

The GENSI framework was applied to integrate transcriptomic data from two hepatocellular carcinoma cell lines, Huh7 and PLC, and NA data. In our study, NAs were limited by the composition of the medium. We cultured both cell lines in six different conditions with various concentrations of glucose and glutamine. We call the NA data from the medium with 25mM glucose and 4mM glutamine concentration “NA1”, 25mM glucose and 0mM glutamine “NA2”, 5mM glucose and 4mM glutamine “NA3”, 5mM glucose and 0mM glutamine “NA4”, 0mM glucose and 4mM glutamine “NA5” and 0mM glucose and 4mM glutamine “NA6” (see Table 1 in Material and Methods). Based on the observed experimental differences in growth rate between the media and cell lines we found a scaling factor that translated RAS and MUR data into the fluxes that matched the observed growth rates.

**Table 1.**
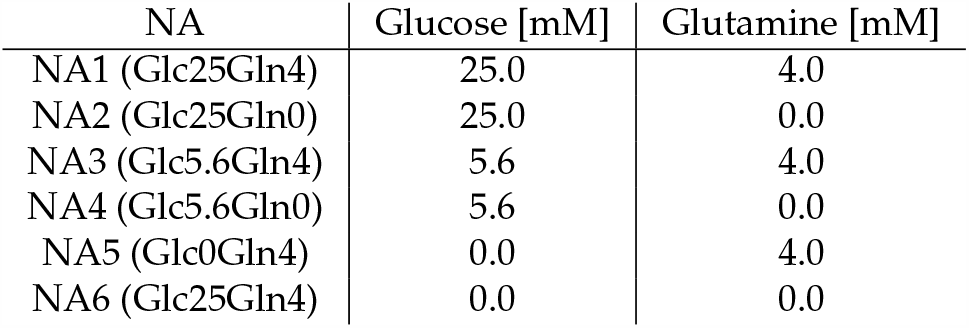
NA data used in experiments in terms of their varying glucose and glutamine concentrations. The NA further contained all components like in standard DMEM medium. The concentrations were verified via HPLC measurements.

| NA | Glucose [mM] | Glutamine [mM] |
| --- | --- | --- |
| NA1 (Glc25Gln4) | 25.0 | 4.0 |
| NA2 (Glc25Gln0) | 25.0 | 0.0 |
| NA3 (Glc5.6Gln4) | 5.6 | 4.0 |
| NA4 (Glc5.6Gln0) | 5.6 | 0.0 |
| NA5 (Glc0Gln4) | 0.0 | 4.0 |
| NA6 (Glc25Gln4) | 0.0 | 0.0 |

We obtained different GEM-RAS-MUR models (see Figure 1) constrained by different combinations of MUR and RAS to investigate the effectiveness of NA and transcriptomic data integration in reducing the metabolic flexibility of the provided solutions. GEM-RAS-MUR is an integrated model obtained by incorporating NA data and RAS data into a GEM model, which is RECON 3D in this work.

### 2.2. Metabolic genes: expression in two hepatoma cell lines

For our two cell lines, the RNA-seq dataset published by Ma et al. [**?**] reports on 17726 genes. We extracted the RNA-seq records for 2232 out of the 2248 metabolic genes that surface both in this data set and in Recon3D (see section Methods). Comparing the mRNA levels (in terms of TPM scores) for the subset of metabolic genes to the genome-wide mRNA levels, we observed that most metabolic genes exhibited a higher than average expression, with a median expression level of ∼ 15 compared to ∼ 1 genome-wide and a mean expression level of ∼ 81 compared to ∼ 39 genome-wide.

We were confronted with a well-known issue with RNA-seq data integration in metabolic models, i.e. zero expression levels, including zeros in mRNAs encoding enzymes catalyzing essential reactions or combinations of enzymes that are essential. Of the 2232 metabolic genes, 262 and 310 had TPM scores equal to zero in Huh7 and PLC respectively. Such zeros could be due to true absence or correspond to technical zeros where the gene had in fact been transcribed but was somehow not measured. Technical reasons may include inefficient cDNA synthesis due to tertiary structure formation, amplification bias, or low sequencing depth. Additionally, zeros may occur due to transcription bursting between somehow synchronized individual cells [59] or too small time-windows of expression. Given that an independently obtained set of microarray data might not suffer from quite the same problems, we assigned to genes with zero TPM scores in the RNA seq analysis, alternative TPM scores that reflected the microarray datasets (see Methods).

Huh7 and PLC being both hepatoma cell lines with a different history of oncogenesis, we further expected them to differ mostly in the expression of oncogenes and perhaps other genes involved in signal transduction or management of the genome, but not much in genes encoding metabolic enzymes. With respect to the metabolic genes, we expected both cell lines to be transcriptionally addicted to the same metabolic Warburg rewiring at the level of transcription. With this expectation, a plot of the expression levels Huh7 versus the expression levels of the same genes in PLC should show all genes close to the diagonal line. Although for the majority of genes the correlation fell close to the diagonal, there were quite a few genes for which the expression levels differed between the two cell lines (Figure 2A).

**Figure 2.**
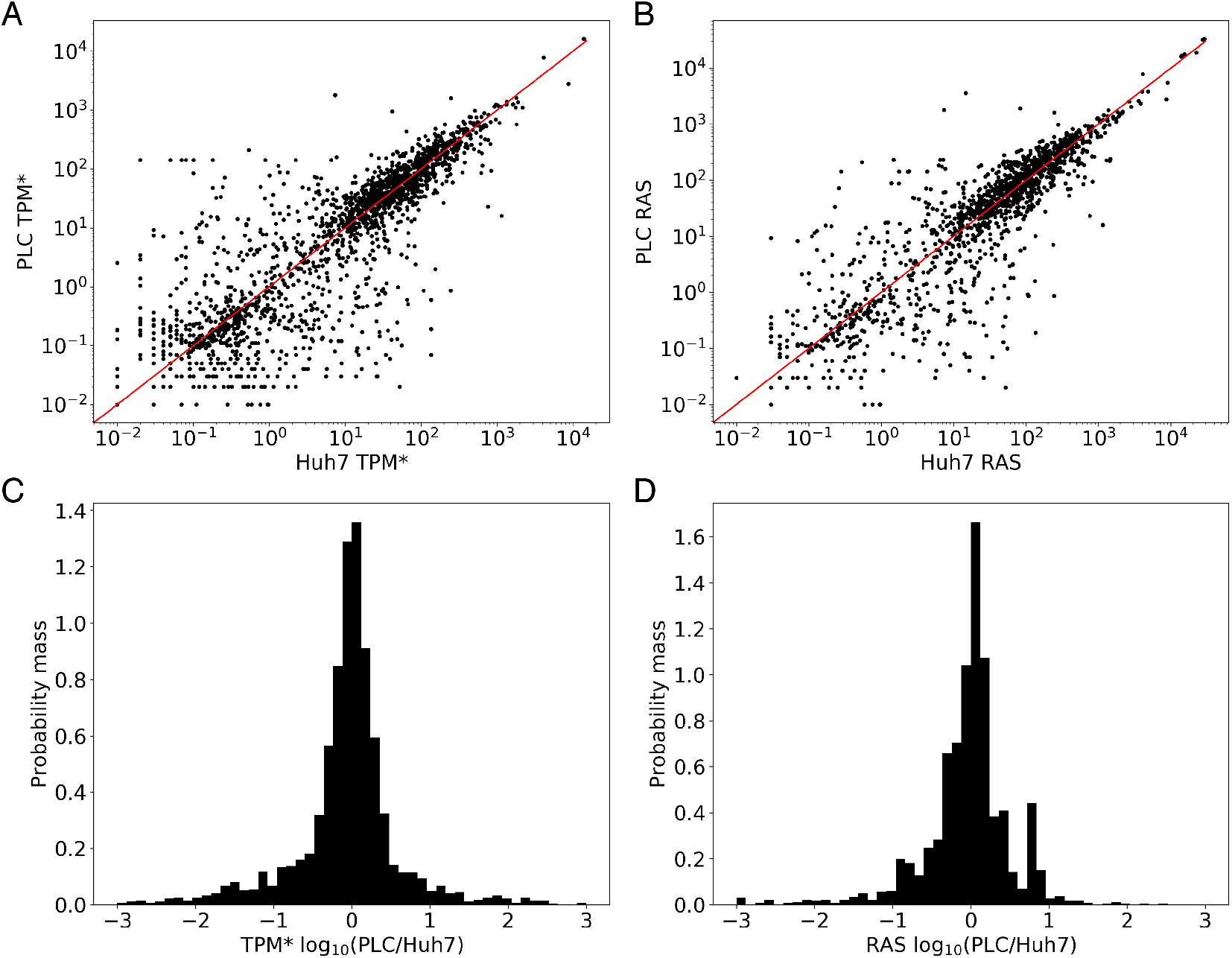
(**A**) Correlation of the mRNA levels (in terms of TPM* scores) between the Huh7 and PLC cell lines for all metabolic genes represented in Recon3D. The line PLC-TPM* = Huh7-TPM* represents theoretical identical scores between the cell lines. (**B**) 2D comparison of the RAS scores (dealing with multi-subunit proteins and isoenzymes) of the Huh7 and PLC cell lines for all metabolic genes represented in Recon3D. (**C**) Histogram of the probability mass function of the log10 of the ratio of the TPM* scores between the two cell lines (PLC relative to Huh7) shown in (**A**). (**D**) Histogram of the probability mass function of the log10 of the ratio of the RAS scores between the two cell lines (PLC relative to Huh7) shown in (**B**).

We first looked at the genes that did correlate: In the RNA-seq dataset (prior to correcting for zeros as discussed above), 186 metabolic genes (∼ 8% of the 2232 genes) did not come with any transcript in either cell line. 215 metabolic genes (∼ 10%) exhibited non-zero TPM values below the genome-wide median (∼ 1 TPM), again in both cell lines. Together, these constituted a common set of 401 metabolic genes that were expressed at a low level. On the high-abundance side, the two cell lines shared 1422 genes (∼ 63%) that were more highly expressed than the median gene. A 623-genes subset of these 1422 (∼ 28%) was commonly expressed above the genome-wide mean (39 TPM). These genes behaved in line with our expectation of metabolic similarity between these cell lines. They came however with a possibly important exception of some 425 genes that were off-diagonal in Figure 2A. 415 metabolic genes (∼ 18%) exhibited a TPM score above the genome-wide median in both cell lines and a differential expression ratio of at least 3 (or below 1/3). The data suggests that our expectation was not quite right: there was an appreciable metabolic difference between the two cell lines.

In accordance with the above, the expression ratio between the two cell lines was still mostly distributed narrowly around 1. The corresponding probability distribution was largely log-normal, but not quite: it had long ‘tails’ on either side, suggesting that a disproportionate number of metabolic genes were much more expressed in Huh7 than in PLC and vice versa (Figure 2C).

### 2.3. Converting RNA-seq data to RAS reduces but does not eliminate metabolic differences between Huh7 and PLC

There are at least two reasons why the different expression of metabolic genes between two cell lines might not affect the activities of the corresponding biochemical reactions. First, the differences in expression level between the two cell lines could be in enzyme subunits that are abundant as compared to other subunits that are equally expressed. Second, the two cell lines may express different proteins (isoenzymes) that catalyze the same reaction. Recon3D dealt with this issue qualitatively through its gene-reaction coupling rules. We used a quantitative version of these rules to assign a Reaction Activity Score. The RAS for a metabolic reaction reflects the expression levels of isoenzymes and components of multi-component complexes that may catalyze that reaction (Materials and Methods, and [28,31,55]). Out of the 10601 reactions in Recon3D, 5938 reactions were assigned a RAS in this manner. For the 2999 reactions catalyzed by single genes, the RAS scores were taken equal to the TPM values. The remaining 4663 reactions are not linked to any genes: they represent so-called ‘exchange reactions’ between the cells’ immediate environment and the outside world, or non-enzyme-catalyzed reactions and transport within the cells or across their membranes. We left the bounds of such geneless reactions unlimited at ±1000.

We then asked whether the metabolic differences we found between the two cell lines would disappear when correcting for these isoenzyme and enzyme subunit issues by assigning reaction activities. In Figure 2B we correlate the RASs between the two cell lines, and panel 1D shows the distribution over the metabolic genes, of the RAS ratios between the two cell lines. The RAS correlation between the two cell lines is only a little stricter than that of the individual mRNAs. The standard deviation in the RAS ratios is 14 compared to a standard deviation of ∼ 177 for the TPM* ratios. The fraction of outliers outside the lognormal distribution of the ratio remains substantial, however. Supplementary Excel Table 3 lists the 92 outlier reactions with a log10 RAS ratio *>* 2 or *<* −2.

### 2.4. Conversion of NA data into MUR

In metabolic networks like RECON3D, all metabolites with defined exchange reactions can be taken up and secreted: i.e. compounds enter and exit the extracellular environment via ‘exchange’ reactions. The GEM is not able to import compounds unless an exchange reaction from the external environment to the inside of the cell is present. To predict the effects of nutritional differences in terms of all components in the medium including the various concentrations of glucose and glutamine, between our six different medium conditions (see Table 1), on the global metabolic behavior of the two cell lines we first converted nutrient availability data into maximum uptake rates (MURs) that describe the rate of maximum possible uptake over time for substances present in the medium. We defined the MUR for the exchange reaction *r*_*j*_ as the absolute value of the difference in the concentration of the substate *s*_*j*_ in the extracellular environment (see methods) over time. In this work, we assume that all amounts of each substance can be uptaken, and therefore the concentration of the substate *s*_*j*_ in time 48h is zero. We equated the upper bound of each uptake reaction to the corresponding MUR, setting the bound to zero if the metabolite was absent from the medium. Export was left unlimited for all metabolites that were allowed to be exported in the default Recon3D map (see Supplementary Excel File S2 for the list of such metabolites). As a consequence of this approach the unit of the uptake fluxes is equal to the unit used for the MUR: i.e. concentration deviated by time. In our study, we set the time to two days (48h) due to experimental conditions.

### 2.5. An FBA-based scaling methodology

In the GENSI framework, we proposed a variant of the approach published by Graudenzia et al. [31]. We set flux bounds of reactions associated with genes proportional to RAS scores, i.e. *b*_*i*_ = *α* · *RAS*_*i*_, where *b*_*i*_ is the flux bound on reaction *i*, is a factor independent of the reaction identity, and *RAS*_*i*_ is the RAS based on the expression levels of an enzyme(s) catalyzing reaction *i*. When reaction *i* is reversible its forward flux is bounded by *b*_*i*_ and its backward flux is bounded by −*b*_*i*_. When reaction *i* is irreversible the flux bound in the impossible direction remains zero and the flux bound in the possible direction is set to *b*_*i*_. Essentially this approach assumes that the *V*_*max*_ of any enzyme is proportional to the corresponding mRNA transcript level. Our variant of the MAREA approach allows us to scale the RAS data, in reference to a specific dataset of NA (see above).

It is a-priori unclear how large the factor should be. There are two possible sources of limitation for the model: the MUR (as represented by maximal uptake rates) and the RAS (consequent to transcriptomics). We here wish to examine the case where the enzyme expression levels begin to impose limitations on the model output. We perform FBA analysis [55] starting with high *α* values, where the enzyme expression levels are not limiting, and maximal biomass flux is determined by substrate concentrations in the medium. Then we decrease *α* and hence all resulting metabolic flux bounds uniformly until we see differential effects on the predicted maximal growth rates across media conditions. Where this occurs significantly, we fix *α*, which thereby becomes a fitted parameter.

We analyzed RAS with the focus on metabolic differences between the cell lines by computing the steady-state flux pattern for the maximal biomass synthesis flux for each medium and for each cell line across a range of values for the factor *α* ∈ [3 × 10^−4^, 1.6] (Figure 3). For either cell line Figure 3A shows that for factor *α* in excess of 0.5, the predicted growth rates differed between media conditions, the ones at 25 mM glucose being about double those at 5.6 mM glucose. Also, dependence on glutamine concentrations was predicted, but this dependence was smaller.

**Figure 3.**
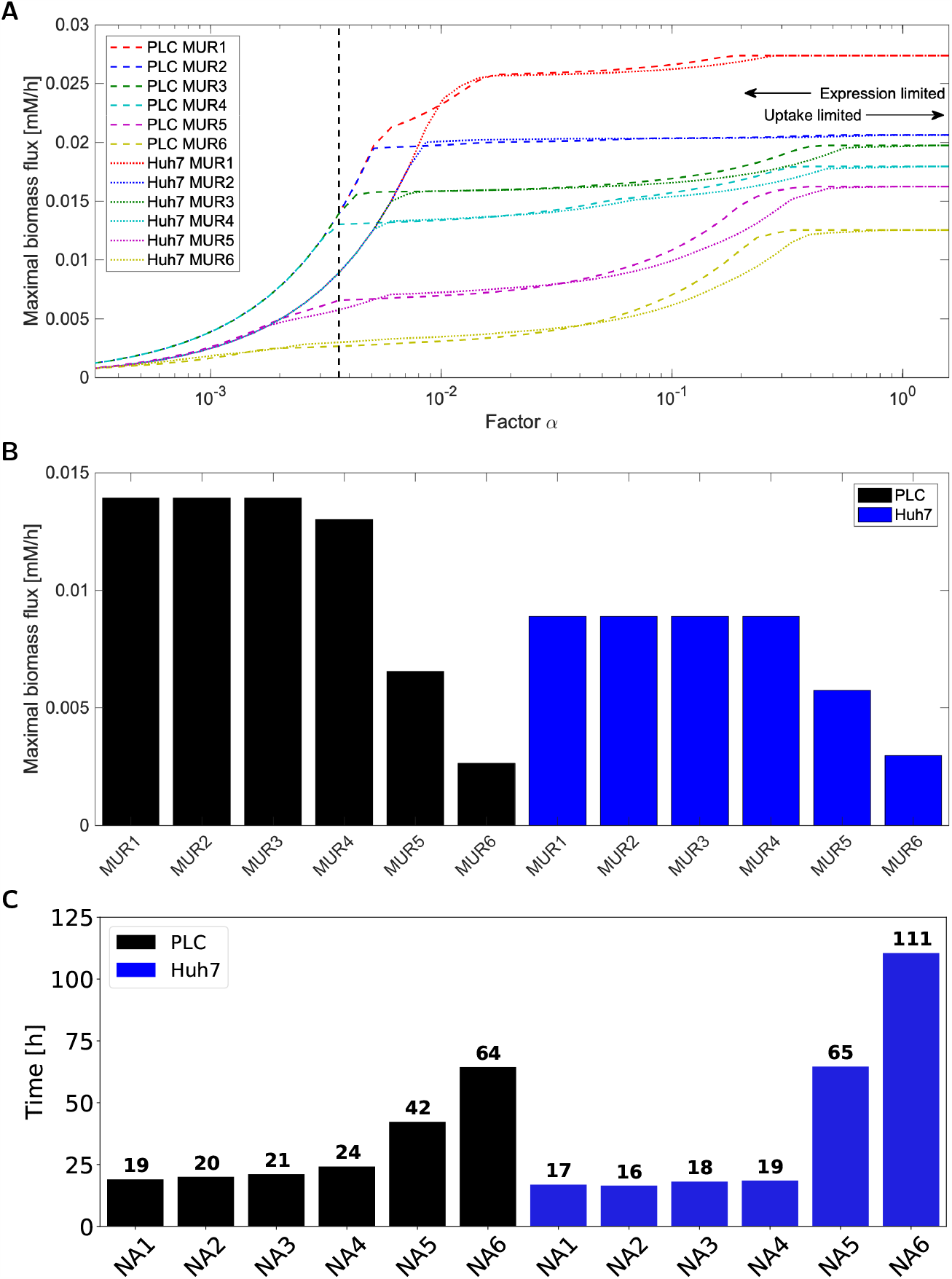
(**A**) Maximal in-silico biomass flux predictions for Huh7 (dashed lines) and PLC (dotted lines) on six MURs versus the factor used to convert RAS to flux bounds. The labels MUR1-6 corresponds to the different medium conditions listed in Table 1 and Table S4 and were effected as proportional uptake (exchange) bounds: Glucose and glutamine were (in mM) 25, 4 (MUR1), 25, 0 (MUR2), 5.6, 4 (MUR3), 5.6, 0 (MUR4), 0, 4 (MUR5) and 0, 0 (MUR6). The dashed black line indicates *α* = 0.0036, the value of the factor *α* that we shall consider more in detail below and at which the maximal biomass flux predictions for MUR1-4 become virtually equal for both cell lines. (**B**) Maximal in-silico biomass flux predictions for PLC and Huh7 across the six GEM-RAS-MUR models. (**C**) Reciprocal of the cells doubling time calculated from a linear equation describing the change in cell number over the time for PLC and Huh7 across the six different media conditions.

When decreasing the factor alpha to below 0.5 the first effects of the transcriptomics start to be noticeable: i.e. growth predictions for the two cell lines start to diverge in some medium conditions. When decreasing the factor *α* to below 0.004, the MUR dependence disappeared for four out of six modeled medium conditions: the same biomass flux was then predicted which was still significantly higher than that for the remaining two medium conditions at zero glucose (5 and 6). Huh7 predictions for MUR 1-4 converged already at higher values for *α* than did the predictions for PLC. Huh7 was predicted to have equal or lower growth rates than PLC across all conditions. MUR 1, followed by MUR 2, was predicted to yield the highest biomass fluxes for the high values of *α*, corresponding (see above) to the absence of gene-expression limitations. This reflects the model’s sensitivity to carbon input for high *α* values since medium 1 and 2 contained the highest levels of glucose. *α* = 0.004, the simulations for MUR 1-4 yielded equal biomass fluxes which were still larger than the predicted fluxes for MUR 5 and 6 which lack glucose. This indicates that by reducing the factor *α*, the model can be made more (high *α*, hence no limitation by low transcription of metabolic genes and thereby limitation by uptake) or less (low *α*, hence strong limitation by low transcription of metabolic genes) sensitive to variation in concentration of the growth substrate.

The fact that MUR 1-4 converges to similar biomass synthesis flux optima for low levels may reflect a shared limiting reaction downstream of (and at a flux bound smaller to the bound of) the exchange reaction the flux bound of which keeps monotonically decreasing with decreasing *α*. In Figure 3B we summarize the predicted maximal biomass fluxes for the factor *α* = 0.1, which is at the transition between limitation by extracellular substrate levels and intracellular expression levels. It shows that reduction of glucose concentration does not decrease maximal biomass flux as we observed in *in vitro* experiments (compare Figure 3B and 3C).

We observe that nontrivial predictions for limitations imposed by medium composition and gene expression can be computed by GENSI method, such as that (i) both in the absence and in the presence of glutamine the growth rate should be independent of glucose concentrations between 5.6 and 25 mM, yet decrease appreciably in the absence of glucose, (ii) the specific growth rate of Huh7 cells is lower than that of PLC cells, (iii) in the absence of glucose, the cells should be able to grow on glutamine, but (iv) growth rate on glutamine alone should be much lower than on glucose alone.

### 2.6. Metabolic flux potential as predicted by Flux Variability Analysis

FBA is oblivious of metabolic regulation other than that it philosophizes about what flux should be optimal for the cell in view of some objective. The transcriptome and extracellular-concentrations informed flux bounds that we here implemented, merely define ranges of the fluxes rather than that they precisely predict the fluxes. Moreover, fluxes through intracellular biochemical reactions are also determined by metabolic regulation [60], i.e. by the concentrations of intracellular metabolites. For the precise predictions of fluxes, one needs fully dynamic models [61]. However, the kinetic information required for this approach is missing for mammalian cells.

We hereby can only predict the ranges of fluxes that are consistent with transcriptome and extracellular nutrient concentrations and this is done here by flux variability analysis (FVA) [62] to test the prediction of the GEM-RAS-MUR models. We performed FVA analysis for allowed exchange reactions while supporting the biomass production rate. For the factor value *α* = 0.0036, indicated by the black vertical line in Figure 3, we analyzed each of the 12 GEM-RAS-MUR models in terms of the possible ranges its production fluxes of lactate, pyruvate, ATP and CO_2_ and its uptake fluxes of glucose, glutamine, oxygen, and phenylalanine. Here we maintained the maximal biomass flux for each specific medium and transcriptome (i.e. the biomass fluxes listed in Figure 3, which differ between the 12 GEM-RAS-MUR models) (Figure 4). The results for lactate and pyruvate production are non-negative since these compounds are not in the growth medium and thus cannot be taken up. In order to avoid thermodynamically infeasible ATP synthesis, the ATP hydrolysis reaction was non-negative by design. The fluxes in Figure 4 for glutamine and CO_2_ can be both negative and positive due to these compounds being present in the medium. If the lower end of its bar in Figure 4 is positive that compound must be produced for the cell to grow at maximal growth rate: it is a primary metabolite. When the upper end of the bar is negative it indicates that the compound needs to be taken up for maximal growth.

**Figure 4.**
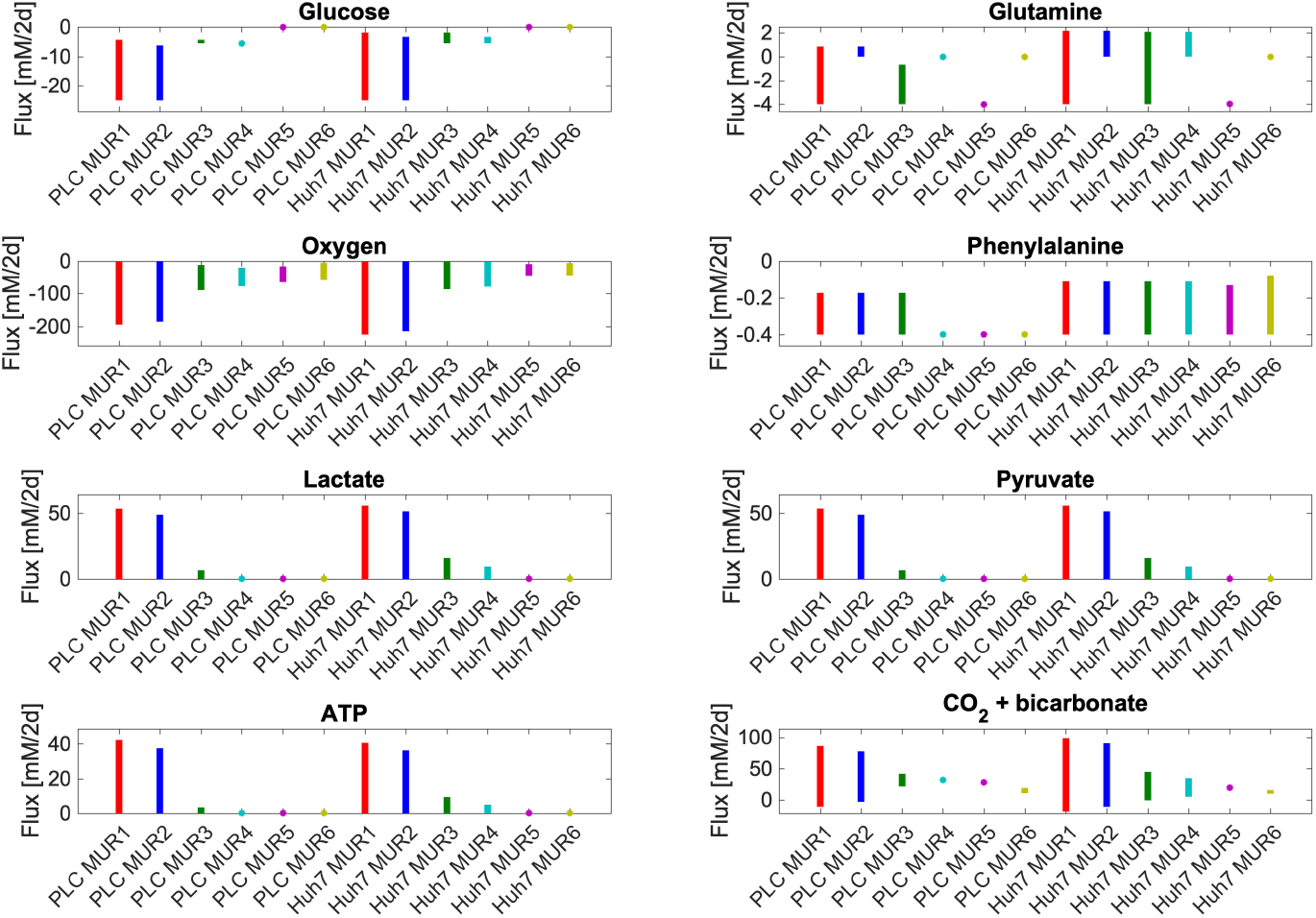
The range of uptake (if negative) / secretion (if positive) fluxes of various metabolites: computed to be consistent with maximal biomass flux and steady-state. Uptake and secretion fluxes of metabolites have units of mM/h. Since we set maximal uptake fluxes equal to medium concentrations these rates differ from reality by some undetermined factor which is identical for all conditions. Minimal and maximal exchange fluxes for each of the compounds lactate, phenylalanine, glutamine, pyruvate, oxygen (O2) and CO2 + bicarbonate and ATP hydrolysis flux (i.e. ATP + H2O => ADP + P*i* + H^+^), were calculated using flux variability analysis [63] while requiring the model to produce the same maximum possible biomass fluxes shown in Figure 3. Maximal glucose and glutamine uptake fluxes had been set to their medium concentrations divided by 2 days (see Table 1). In these calculations, ATP is treated differently from the others. For the others, the reaction was and remained present and potentially carrying varying flux when any of the yet other fluxes was manipulated in the FVA. For the ATP, the ATPase reaction was absent (no growth rate-independent maintenance, therefore) when doing FVA for any of the others and only present when ATP synthesis was manipulated by forcing flux through an ATP hydrolyzing added reaction. For visibility, we used starred markers to indicate small flux ranges. These markers are typically located at minimal or maximal flux boundaries: e.g. zero glucose uptake in NA5 and NA6 or maximal phenylalanine uptake in PLC NA4. Uptake and secretion fluxes of metabolites have units of mM/2d.

In Figure 4 we see that in the media where glucose is present (NA1-NA4) some glucose uptake is essential for attaining the maximal growth rate. In NA5-NA6 glucose uptake is always zero due to its absence from the medium. Phenylalanine, an essential amino acid, functions as a positive control. Its uptake proves indeed essential in all media for both cell lines as expected. In most media, its uptake rate can vary from the amount of phenylalanine in biomass to 0.4, the rate at which it can be used to provide nitrogen to other parts of anabolism. This maximum uptake rate of 0.4 [mM/2d] is the same for all media and both cell lines, reflecting that this corresponds to its concentration in the media. In the absence of glutamine and in the presence of low glucose (NA4), PLC needs to make full use of this phenylalanine in order to achieve its maximum growth rate, but Huh7 cells could still vary the amount of phenylalanine used whilst attaining the same growth rate. Maximum lactate, pyruvate, and ATP production capabilities track the total amount of carbon in the medium within each cell line, with subtle differences between the two cell lines. The gene expression levels appear to be consistent with shifting to virtually complete metabolism of glucose and glutamine to lactate whilst maintaining the maximum growth rate. At the same biomass production flux, lactate efflux could also be as high as 55 mM/2d, roughly corresponding to 2 lactate per maximum glucose consumed plus one lactate per maximal glutamine consumed. However, in all models, the lactate secretion can also be zero while maintaining maximal growth. This shows that glucose conversion to lactate can vary greatly, and may also reflect that in our models the cells can produce and secrete other compounds such as pyruvate. The *in silico* cells are not addicted to the Warburg effect.

Glutamine uptake is only essential for both PLC and Huh7 in medium 5 and for PLC only in medium 3. In medium 3 for PLC it is then required at a very low amount to achieve optimal growth (as indicated by the upper end of the bar) whereas in medium 5 in both cell lines the maximal uptake bound has to be hit to achieve maximal growth (see the markers at −4). Because (*in silico*) the maximum growth rate of Huh7 is lower it has the luxury of producing glutamine from glucose whilst growing maximally in M3 whereas this is not possible for PLC. Both cell lines may produce glutamine in M1 and M2 owing to the excess glucose in those media. We conclude that the model cells are insensitive to glutamine concentrations in the medium in the presence of high glucose but glutamine-sensitive in the absence of glucose.

In all models, a small amount of oxygen must be taken up. CO_2_ (either as CO_2_ or as bicarbonate) may either be produced or taken up in M1-M2 for both cell lines and M3 for Huh7 and may only be produced in M3 for PLC and M4-M6 for both cell lines. CO_2_ uptake might have to do with the reversal of the isocitrate dehydrogenase reaction, which produces isocitrate as a substrate for ATP citrate lyase producing cytosolic acetyl CoA for lipid and cholesterol synthesis [27]. For conditions where CO_2_ or bicarbonate production is required, this may point to oxidative phosphorylation being required for maximal biomass production. Oxygen uptake was essential even in conditions where oxidative phosphorylation (interpreted as CO_2_ production) seems not to be required. In these cases, oxygen uptake may be necessary for the synthesis of tyrosine, cholesterol, and other lipids that are part of the biomass definition and absent from our growth media. We checked that removing cholesterol from the biomass equation reduced the need for oxygen, but it did not remove it.

The possibility to grow at a maximum rate in the absence of CO_2_ production in some conditions highlights the possibility for cells to obtain all the Gibbs energy they need for maximal growth only from the conversion of glucose to lactate. This may underlie the selection of the Warburg effect by a-social cells. It does not quite correspond to the Warburg and WarburQ effects, however: the *in silico* cells are not addicted to the absence of respiration, as they can still respire all this substrate whilst growing at the same rate. In PLC cells, but not in Huh7 cells, the maximal growth rate in low glucose medium without glutamine (MUR4) does require oxidative phosphorylation, consistent with the glutamine to lactate pathway elucidated by Damiani et al [22]. These and other apparently minor differences between cell lines in our FVA results are of interest, as they suggest that drugs, in this case, ones that inhibit respiration, should be effective against some cancer cells and not others, also depending on extracellular metabolic conditions.

We further explored this by plotting some essential reactions for respiration to occur analogously to Figure 4 in terms of their minimal and maximal possible flux allowed while maintaining maximal biomass flux for the medium and cell type specified (Figure 5). In media, with glucose, the maximum growth rate does not require flux through cytochrome oxidase and oxidative phosphorylation with the exception of NA4 for PLC. Figure 5 suggests that for PLC respiration in terms of flux through cytochrome oxidase is required for maintaining the maximal biomass flux in media 4-6 whereas for Huh7 this is required for media 5 and 6: rather than a range of fluxes, a precise non-zero flux magnitude is required. This suggests that only in those cases of limiting metabolic substrate, the maximal growth rate depends strictly on ATP produced by oxidative phosphorylation. This is in full agreement with the interpretation of the CO_2_ + bicarbonate panel in Figure 4. In all other cases, respiration is optional for maximum biomass synthesis flux, suggesting that the cells can obtain their ATP from other processes including aerobic glycolysis. To maintain their maximum growth rate at the 5.6 mM glucose concentration, they do need to use virtually all that glucose, however (Figure 4).

**Figure 5.**
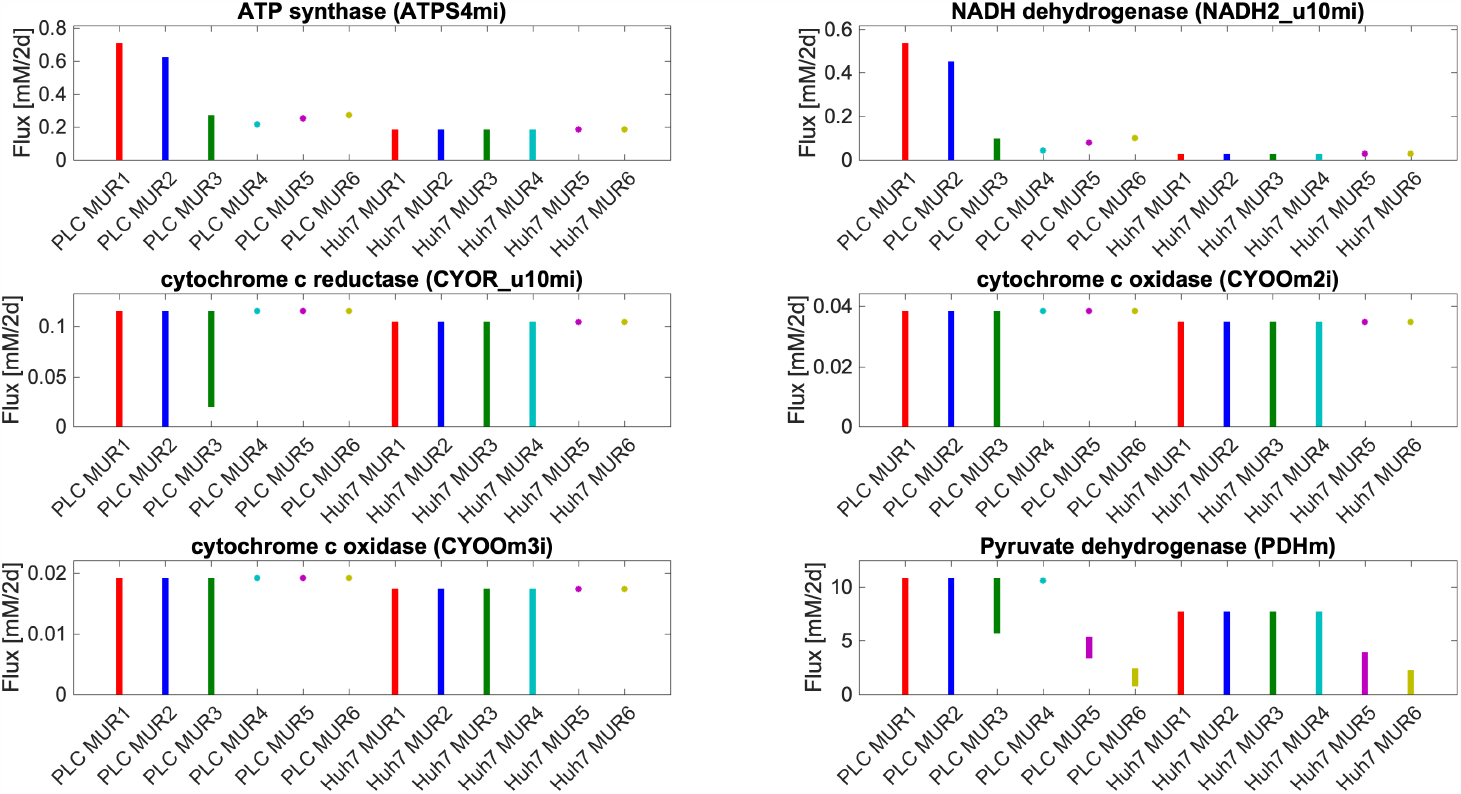
Range of allowed flux values through various reactions related to mitochondrial oxidative phosphorylation while maintaining the maximal biomass flux for the medium condition and cell type specified on the abscissa. See Table S3 for the detailed reactions. Uptake and secretion fluxes of metabolites have units of mM/2d.

Because of its assumptions of maintenance of maximum growth rate and the full capability of the network to allow for various fluxes, flux variability analysis makes few predictions that may be put to the test in this study. Exceptions are (i) both cell lines should be capable of consuming the 5.6 or 25 mM glucose offered to them, (ii) they are not addicted to a 100% aerobic glycolysis, but can reduce lactate production without giving up their maximum growth rate, (iii) at glucose concentrations around 5 mM they would make use of all that glucose to grow maximally, (iv) They should be capable of catabolizing glutamine both in the absence and presence of glucose.

### 2.7. Experimental verification

To verify whether GEM-RAS-MUR models obtained via GENSI give reliable predictions, we measured the concentration of the crucial components in the medium during culturing. We observed a decrease in the concentration of glucose with time such that some 5 mM was consumed by the PLC cells (Figure 6B). In the PLC cultures that started with 5.6 mM glucose, this resulted in almost full glucose depletion after the first day and night. In the case of the Huh7 cell line, the consumption of glucose was higher in rich glucose medium than in low glucose medium reaching some 8.5 mM and 5 mM, respectively. The further velocity of glucose consumption for Huh7 was the same, while in the case of PLC cells, after 24 h of culturing, stabilization appeared and lasted for the next 20 hours (Figure 6B and 6C). We did not observe a difference in glucose consumption between media with and without glutamine. With respect to glutamine consumption by the cells, we found that the level of glutamine did not change maintaining the 4.5 mM level for media with glutamine (that contained originally 4 mM) and 0.5 mM without.

**Figure 6.**
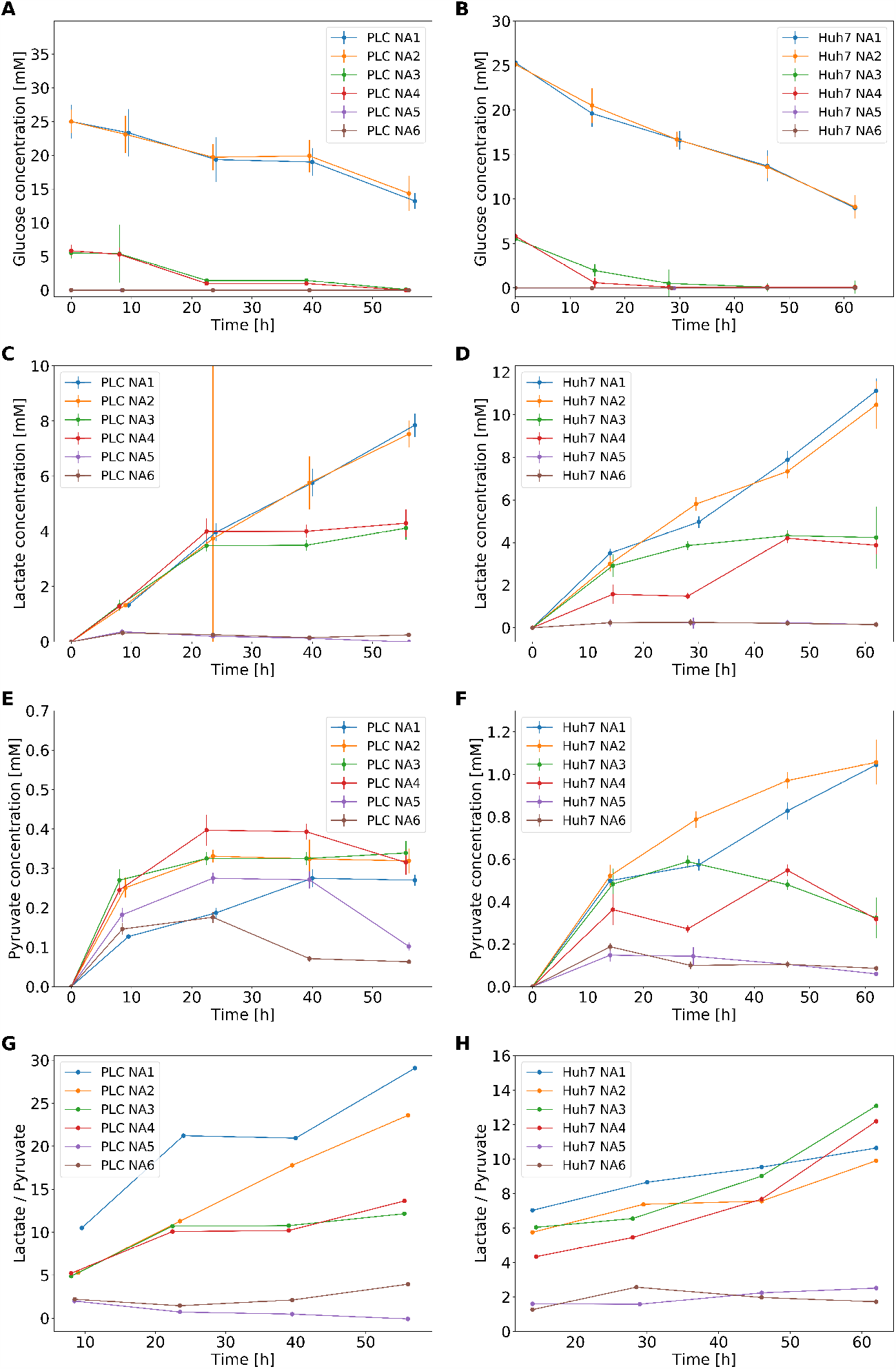
Metabolic performance as a function of time for Huh7 and PLC cells in six different media. Glucose consumption by PLC (**A**) and Huh7 (**B**) in four different media. In media M5 and M6 glucose was not detected. Lactate production by PLC (**C**) and Huh7 (**D**) cells. Pyruvate secretion by PLC (**E**) and Huh7 (**F**) cells. Ratio of extracellular lactate to pyruvate in PLC (**G**) and Huh7 (**H**) cells starting from the second time point measured in panels **D**-**G**. In all six media the concentration of lactate and pyruvate before the experiments was 0 mM.

We additionally measured the production of lactate and pyruvate by cells using the same media conditions as above. In the case of the PLC cell line, the production of lactate was the same during the first 24 hours for NA1-NA4, the lactate concentration reaching some 5 mM at the end of this time period (Figure C and D). After 55 hours the concentration of lactate was twice higher in samples derived from the glucose-rich NA1 and NA2 (8.5 mM) than in samples taken from the low-glucose (5.6 mM) media NA3 and NA4, ostensibly because after 24 h the glucose had run out. In media low in glucose (NA3 and NA4), slightly over 40% of the glucose was consumed during the experiment was transformed to lactate, while for M1 and M2 this was a bit less, i.e. 35%. For the Huh7 cells, the trends were similar in the case of NA1 - NA3. In the case of NA4, such production was achieved only after 45 hours. In NA1 and NA2 lactate increased linearly with time attaining 10 mM at the end of the experiment.

Pyruvate production was quite small for PLC cells, from 0.15 mM (NA1 and NA6) to 0.4 mM (NA4) during the first 24 h. Pyruvate production was higher for media without glutamine (NA2, NA4, and NA6 vs. NA1, NA3, and NA5). For PLC cells, the lactate/pyruvate ratio (L/P) showed a positive relationship with the glucose consumption, being highest for NA1, and lowest for NA5. For Huh7 cells, the time-series for pyruvate (Figure 6F) look similar to those that show lactate secretion (Figure 6D). On NA1-4 the ratio of lactate to pyruvate changed over time from 6 to 10 whereas it stayed roughly constant for NA5 and NA6.

Consistent with the hypothesis that both cell lines exhibited a Warburg effect, they produced an amount of lactate proportionate to the amount of glucose that was present initially. Apparent differences merely derived from the glucose running out after some 30 h in the experiment starting at only 5.6 mM of glucose. Only part of the glucose was transformed to lactate, around ∼ 43% for low glucose media and perhaps a little as ∼ 33% in glucose-rich media. Thus, lactate production corresponded to only part of what might have been expected for full glucose conversion to lactate. This deficit in lactate secretion can be explained by the utilization of glucose for oxidative phosphorylation and as a source of carbon for the new biomass. Consistently, we did not observe any difference between NA1 and NA2 for both cell lines nor did we observe such differences between NA3 and NA4 for PLC; for Huh7 small deviations were noticed. For the two cell lines we examined, this proves the independence of lactate production from glutamine access.

## 3. Discussion

In this work, we follow recently exploring interest on the impact of the diet on cancer cell metabolism and progression. We explore the idea that the utilization of nutrients by cancer cells is determined not only by cancer genetics but also by the metabolic environment.

To investigate whether cancer progression may be mediated through changes in the access to nutrients, genetics and the availability of nutrients have to be employed simultaneously. Genome-scale models that integrate all known metabolic reactions occurring in an organism into a single map give us such an opportunity. The gene-reaction coupling rule and exchange reactions allow for the integration of gene expression and nutrient availability data, respectively. Here, we aim to provide an FBA-based framework, named GENSI, for studying the influence of the nutrient availability on the rate of proliferation and metabolic phenotype based on transcriptomic and NA data. Our method could be applied to the study of every cancer type and therefore could lead to a better understanding of how diet impacts cancer cell metabolism and identifying how different cancer types respond to different nutrients composition. GENSI translates the relative importance of gene expression and nutrient availability into the fluxes and generates high-quality GEM-based specific models (GEM-RAS-MUR) that can be used to predict metabolic changes upon nutrition shift by flux balance analysis.

Further, we applied the proposed method to study the influence of the glucose and glutamine availability on their consumption, some metabolic signatures, and the growth rate of the hepatocellular carcinoma cell lines PLC and six for Huh7. We, therefore, selected cell types that do exhibit the Warburg effect clearly. We used transcriptomic data from the cell population in its entirely and the models are a representation thereof. Single-cell transcriptomic studies suggest that cancer cell populations are heterogeneous, but single-cell metabolomics does not yet enable us to examine the consequences of metabolism. Thus, we assumed in the model that the cell population was homogeneous.

Our findings offer support for the predictive potential of genome-scale metabolic maps together with transcriptomic data sets and nutrient availability data. The predictions address the carbon and energy metabolism of cancer cells. Because much of this metabolism is essential for cell survival, the potential may translate to new drug targets in a long-neglected area of drug discovery. Indeed, we have shown that dependencies on nutrient availability and gene expression can be computed and that the results are then relevant enough to be compared with experimental work.

Notwithstanding these successes, our methodology comes with a number of issues. One of these relates to the translation of the expression level information to flux. In contrast to several methods developed to extract context-specific models [32,33,35] that focused on threshold selection with exception of some essential metabolic functions that are needed for cell growth, our method used a linear relationship between flux bounds and the transcriptome. Such a linear approach was first proposed by Colijn et al. [28] and applied in the MaREA framework by Graudenzi et al. [31], where it was shown to be of use in comparing the metabolism of samples in distinct subgroups. The approach assumes that the *V*_*max*_ of a reaction is proportional to the level of the mRNA encoding the enzyme, with a proportionality constant equal for all reactions. It thereby neglects differential translation and posttranslational regulation, and assumes all *k*_*cat*_ to be equal. Furthermore, we assume that the cell lines in our experiments have not significantly evolved compared to those used in the transcriptomics datasets [**?**]. We did not take into consideration epigenetic and gene-expression changes that could occur during culturing. In addition, our approach equated MUR data (that we obtained based on NA data) with exchange reaction lower bound, again treating all compounds equally. The limitations of this step include the failure to take into consideration the kinetics and expression levels of transporters. What is also peculiar in our approach is the arbitrary magnitude of the ratio of the proportionality constant relating RAS level to enzyme bound to the proportionality constant relating MUR to exchange bound. We mediated this problem by setting the factor *α* to a value that enabled the models to shift from limitation by nutrient availability to limitation by expression level.

*In silico* experiments on GEM-RAS-MUR models showed that the specific growth rate of Huh7 cells is lower than that of PLC cells, which was also observed during *in vitro* experiments on these cell lines. Given the positive correlation between the *in silico* biomass flux and the experimentally determined growth rate, we speculated that the *in silico* results pertaining to the range of uptake and secretion fluxes of various metabolites might also correlate with experimental results. Modeling showed that in the absence and in the presence of glutamine the growth rate should be independent of glucose concentrations between 5.6 and 25 mM. Also, experimentally the growth rate reached the same level for four nutrient availability (NA1-NA4) that contained 5.6 and 25mM of glucose for both HCC cell lines. Moreover, both approaches, computational and experimental, showed that in the absence of glucose, the cells should be able to grow on glutamine but with a much lower growth rate than on glucose alone.

According to the flux variability analysis predictions, both cell lines should have a higher potential; they should be capable of consuming the 5.6 or 25mM glucose offered to them. Yet, our *in vitro* observations indicated that PLC cells confronted with the 25mM did not consume more than the around 5mM during the first 24 hours. They did not make use of that full potential. We conclude that there is a maximum amount of glucose per unit time that the cells ‘wish to’ handle at glucose levels above a few mM, because other issues than energy and carbon may limit the cells’ growth. We use the term ‘wish to’ to indicate that this may be an issue of metabolic regulation: the gene expression would allow for higher glucose uptake fluxes.

The cells that had consumed in the first 24h some 5mM of the 5.6mM glucose they had been incubated with, continued to grow for the next 24 hours at virtually the same growth rate; they must have done this at a much-reduced glucose consumption rate. Since they also stopped producing lactate, we suspect that the small amount (approximately 0.5mM) of glucose left provided the cells with sufficient ATP to drive their continued growth. The cells may have reverted from lactate production to respiration with its more than 15-fold higher ATP yield. This highlights that there may be a limitation to these cells’ addiction to lactate production: these cancer cells can shift to glucose oxidation. We observed a slightly different effect in the case of the Huh7 cell line: the consumption of glucose was almost two times higher in rich glucose medium (25 mM) than low on (5mM) during the first 24 hours. *In silico* analyses predicted and *in vitro* experiments confirmed that both cell lines, at glucose concentrations around 5mM, made use of all available glucose to grow maximally. They also showed that upon glucose depletion and if asked to grow at the maximal growth rate, the cells should shift to glucose respiration (Figure 4 and 5).

Significant progress has been made toward developing the tool to study how diet and nutrition affect cancer progression. We obtained GEM-RAS-MUR models for the hepatocarcinoma cell lines and used them to predict cell growth and metabolite consumption/production. Our methodology could be applied to different types of cancer to investigate how cells respond to the diet and how to mediate these responses. Potential future applications of the GEM-RAS-MUR model would be to predict cell addictions and metabolic requirements in order to modulate them for dietary aid in cancer treatment. Moreover, GENSI methodology is not limited to cancer research, it could be applied to study the metabolism of any healthy and/or sick cells to study how diet and nutrition affect metabolic phenotype, proliferation, and functions of the cell.

## 4. Materials and Methods

### Creation of specific GEM

#### Inputs

##### GEM Model

The most comprehensive genome-scale model of human metabolism RECON3D [38] that includes information on 3,288 open reading frames that encode metabolic enzymes catalyzing 13,543 reactions on 4,140 unique metabolites, was used in this study.

##### RNA-seq data

RNA sequencing data for both cell lines was obtained through NCBI’s GEO database. Specifically, the RNA-seq dataset from [57] was accessed from: https://www.ncbi.nlm.nih.gov/geo/query/acc.cgi?acc=GSE86602 on November 7th 2018. The RNA-seq dataset pertains to cells grown in DMEM containing high glucose (Gibco BRL, Grand Island, NY), 10% heat-inactivated fetal bovine serum (Gibco BRL), 100 mg/mL penicillin G, and 50 µg/mL streptomycin (Gibco BRL) at 37°C in a humidified atmosphere containing 5% CO_2_ [57]. The RNA-seq data records transcript levels in terms of a TP(K)M value (Transcripts Per Kilobase Million): i.e., read count normalized by gene length in kilobases (RPK) and divided by 1 million. Microarray transcriptomics data. We obtained microarray transcriptomics data from the MERAV database [58] (http://merav.wi.mit.edu/) for both the Huh7 and PLC/PRF/5 cell lines. There was data available from two experiments for Huh7 and one for PLC/PRF/5. All MERAV microarray datasets were renormalized together [58]. For the two Huh7 experimental datasets we averaged the signal per gene between the two experiments.

### Nutrient availability data

The formulation of the media used in cell culturing was used as Nutrient availability data. The standard Dulbecco’s Modified Eagle Medium (DMEM) with different concentration of glucose and glutamine was used (see Table 1).

### Preprocessing

#### Preparing the model

During preliminary calculations, we observed that with the default Recon3D model and constraints applied in this study, biomass synthesis was not possible. We traced the problem back to the triglyceride synthesis pathway where in the default Recon3D version a ‘source’ reaction for triglycerides is present (which allows influx into the cell independent of the presence of triglycerides in the medium; it may be noted that in Recon3D such a source reaction is called a sink reaction, by virtue of sign notation; uptake fluxes are positive, effluxes are positive [55]). Indeed, when temporarily reactivating the triglyceride source reaction (or equivalently activating the triglyceride exchange reaction and adding triglyceride to the medium) a biomass synthesis became possible in all conditions.

Biochemically, triglycerides are synthesized starting from glycerol-3-phosphate and various lipid tails esterified to CoA. Each of these lipid tails can assume any of the three positions in the triglyceride molecule. We observed that without adding triglyceride uptake (or a ditto source) to the metabolic map, the first intermediate in the pathway (lysophosphatidic acid; the monoglyceride with a phosphate on the 3 position) could not be net-produced (see the network diagram in Figure S1 and S2). Inspection of the network revealed that this was due to the need for net input of ‘Rtotal’, ‘Rtotal2’ and ‘Rtotal3’ groups in this pathway for which there exists no synthesis reaction in Recon3D. This problem affects biomass flux because multiple metabolites downstream of the triglyceride synthesis pathway (e.g. Phosphatidylcholine, Phosphatidylserine, and Phosphatidylethanolamine, see Table S2), or in branches of it, are explicit components of the biomass used in Recon3D. Recon3D does have the potential to make Stearoyl-CoA, Palmityl-CoA, Oleoyl-CoA and Octadecadienoyl-CoA, but lacks the reactions to associate these with the glycerol moiety: it instead associates Rtotal, Rtotal2 and Rtotal3 to the glycerol moiety, the numbers referring to the position they take in the resulting triglyceride molecule. Recon3D worked around the ensuing problem of lack of biomass synthesis by adding a source for triglycerides. We removed such dei ex machina by forbidding source reactions and thereby came across this problem. We solved it by equating Rtotal, Rtotal2 and Rtotal3 species in Recon3D to a single pool Rtotal and by adding a pooling reaction for lipid tails:

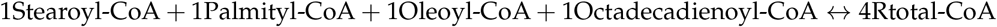

This apparent synthesis reaction merely reflects that the four acyl groups mentioned may be considered a single pool, in the sense that they can largely substitute for each other as lipid tails in biomass. With these two adjustments we have essentially reverted the Recon3D model back to how these reactions were annotated in Recon 2.2 [63,64] but with a different Rtotal synthesis reaction. In Recon2.2 Rtotal synthesis was only possible from Palmityl-CoA and Palmitoleyl-CoA in separate reactions. From Icosanoyl-CoA and Stearoyl-CoA Rtotal2CoA and Rtotal3CoA respectively, could be synthesized but these metabolites were not connected to anything else in the network. Our grouping of the synthesis of all Rtotal into a single reaction, has the advantage that a ratio may be imposed when this is known experimentally. We here took this ratio to be 1:1:1:1 This workaround allowed Recon3D to sustain a positive biomass flux on the medium discussed above. An alternative in which all Rtotal-CoA could be synthesized from any of the four above CoA esters was not here entertained. The resulting ‘patched’ version of Recon3D is available in the supplementary code and model archive and a list of its reactions and metabolites can be seen in Supplementary Excel Table 2.

#### Conversion RNA-seq to RAS

First, we converted the entrez gene identifiers in Recon3D to their gene symbols (e.g. 8639.1 was converted to AOC3), using the mygene module in Python (https://pypi.org/project/mygene/), in order to match them to genes represented in the transcriptomics datasets. Out of the 2248 genes in Recon3D, we were unable to match 103 to the dataset on the basis of their gene symbol alone. We then additionally searched the dataset for known gene aliases and this led us to identifying an additional 87 transcripts. 16 genes of Recon3D (see Table S1) we could not identify. These included the three mitochondrial genes encoding the 3 subunits of cytochrome c oxidase. We artificially assigned all these 16 genes a TPM score equal to the maximum TPM score of the genes we could identify for each cell line. Since we do not know whether or not these genes are expressed, we did not want to artificially block them in our model analyses. Setting them to the maximal observed TPM value guarantees that they will not be limiting in any of our analyses.

The existing RNA-sequencing methodology suffers from so-called zero-inflation [65], i.e. the lack of transcripts for genes that are in fact expressed. For data integration this is problematic since a single zero may block an entire pathway. For our dataset we did indeed observe this problem. For example, in the RNA-seq data the serinepalmitoyltransferase-long-chain-base-subunit-3 gene (SPTLC3) which catalyzes the reaction synthesizing 3-dehydrosphinganine (SERPT), has a TPM of zero, and blocking this reaction (after imposing our changes to Recon3D as discussed above) singlehandedly prevents biomass synthesis. To bypass this problem, we used a microarray dataset and calculated the ratio R of the microarray intensity divided by the genome-wide median for each gene that came associated with a TPM score of zero in the RNA-seq dataset. Then we updated such genes’ TPM scores and set them equal to the calculated ratio R for the microarray dataset multiplied by the genome-wide median in the RNA-seq dataset. Below we will therefore refer to this set of zero-adjusted TPM scores as TPM*. Because the microarray dataset had scores for all genes, this removed all zeros from the dataset. We here neglected any transcriptome difference between the cell lines and experimental conditions used for the microarray and RNA-seq experiments and we assumed that the microarray data were quantitative also at low gene expression.

Recon3D contains an annotated gene-reaction coupling rule for each reaction. Using AND and OR logic this rule specifies which genes encode proteins that may help catalyze that reaction. The AND logic may be used to indicate proteins that consist of more than one subunit, or protein complexes that catalyze reactions whereas the OR logic may be used to indicate isoenzymes or alternative configurations of the protein complexes. When integrating the transcriptomics data into the map we turned these Boolean gene-reaction coupling rules into quantitative rules. Here we were inspired by MaREA methodology [31] and the E-flux approach [28]. MaREA had been developed to compare reaction activities between patients rather than between cell lines by assigning a quantitative so-called RAS to each reaction based on gene expression levels for all proteins that might be involved in the catalysis of that reaction also turning the above Boolean rules into quantitative activities. In order to do this, one needs to know the levels of the corresponding proteins and protein subunits in the cell of interest. We used the mRNA levels as a first approximation to the corresponding protein levels, assuming that the two were proportional. We assigned to each reaction a RAS by summing over isoenzymes (for OR logic) and taking minima of subunits of a complex (for AND logic) the TPM scores for the genes coupled to each reaction. In this way, isoenzymes are thought to contribute additively to the activity of a reaction whereas lack of even one subunit of an enzyme complex can linearly bring down a reaction’s activity [31,56].

### Conversion NA to MUR

Once a network reconstruction is converted to a mathematical format, the inputs to the system must be defined by adding consideration of the extracellular environment. In a metabolic network like RECON3D, compounds enter and exit the extracellular environment via exchange reactions. Exchange refers to the net consumption or production of a metabolite and occurs between the extracellular compartment and the outside. In our GENSI approach the GEM is able to import only compounds that are available for cells. We introduced the Maximum Uptake Rate value as the rate of the maximum possible uptake over the time for each substance available to the model. We define a *MUR*_*j*_ for each exchange reaction *j* based on NA data for each condition as the absolute value of the difference in concentration of the substate *s*_*ex*_ in the extracellular environment over the time *t* as follows:

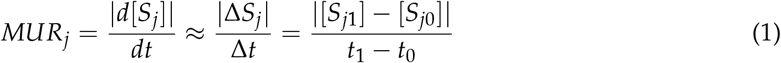

where *S*_*j*_ denotes the concentration of the extracellular substance *j* at timepoint 1 and 0. *MUR* has the units of mol/L/h. In this study, *S*_*j*1_ is zero due to the assumption that the full amounts of each substance could be taken up.

In the next step *MUR*_*j*_ is converted to a lower bound *l*_*j*_ of exchange reaction *r*_*ex*_ as follows:

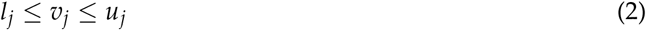

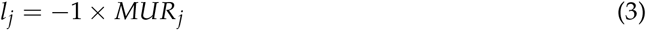

where *v*_*j*_ is the flux through exchange reaction *j, l*_*j*_ and *u*_*j*_ are lower and upper bounds on this flux, respectively, *MUR*_*j*_ is a Maximum Uptake Rate that is defined as the rate of the maximum possible uptake over time for substance *j* that is available to the model.

If the concentration of a substrate supplied to the cells is zero, then the bound (=upper limit) on inward exchange flux is zero (*l*_*j*_ = 0). This does not mean that the transport reaction of that metabolite is blocked (has bounds of zero) but it does mean that that the transport flux will be zero due to lack of substrate. If that extracellular concentration were to go up (by increasing the exchange reaction bound), the metabolite could be imported and transport flux would be possible.

We blocked all other uptake of metabolites so that only those compounds listed in Table 2 were allowed to be exchanged in. We did not alter the Recon3D default choices of which metabolites may be net produced by the in-silico cell. This left 1559 metabolites which are allowed to be net-produced by the cell (Supplementary Excel Table 1). Recon3D also contains various so-called sink and demand reactions which serve as sources and sinks for certain metabolites allowing them to bypass the regular mass-balance. We blocked all such reactions.

We defined NA based on composition on the media used in experiments, we verified the concentration by HPLC after adding serum. All uptake reactions and their maximal uptake rates were taken the same across the six conditions with the exceptions of glutamine and glucose which were varied as specified in Table 1 and Table S4 in Supplementary Material.

### Simulations

In all simulations in this work, we apply the computational technique of FBA [55] and FVA [62] to the human genome-wide metabolic map Recon3D [38] using the COBRA toolbox in MATLAB and Python [66,67]. FBA entails the following linear programming problem:

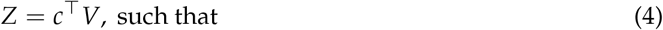

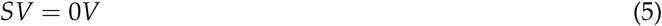

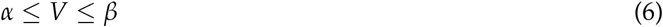

where *S* is the stoichiometry matrix indicating how many molecules of each metabolite are produced or consumed in each reaction, *V* is the vector of fluxes through all reactions including exchange reactions with the environment of the system considered, *α* and *β* are the vectors of lower and upper bounds on these fluxes, and *c* is a vector of weights generating the linear combination of fluxes that constitutes the objective function *Z*. A flux distribution resulting from FBA therefore satisfies the requirements that each metabolite is produced at the same rate as it is consumed, that the flux boundaries are not exceeded and that the flux distribution maximizes (or minimizes) the objective function *Z*.

Whereas FVA:

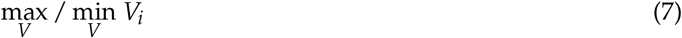

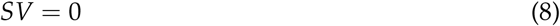

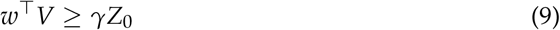

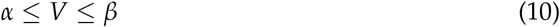

where *Z*_0_ = *w*^⊤^*V*_0_ is the optimal solution to the FBA problem with biomass reaction as the objective function, *w*^⊤^ represents the biomass objective (vector of weights generating the linear combination of fluxes that constitutes the objective function *Z*), *V* is the vector of fluxes through all reactions, *γ* is a parameter that controls whether the analysis is done w.r.t. suboptimal network states (0 ≤ *γ <* 1) or to the optimal state (*γ* = 1).

### Data and source code availability

All Python and MATLAB code, Jupyter Notebooks and raw data files are available as part of a Github repository https://github.com/ThierryMondeel/HCC_flux_balance_analysis.

### *In vitro* experiments

#### Cell culture

The cells were cultured in Dulbecco’s modified Eagle’s media (DMEM; GibcoBRL, Grand Island, NY, USA) at various initial concentrations of glucose and glutamine (M1-M6, table 1 and 2), supplemented with 10% fetal bovine serum (GibcoBRL, Grand Island, NY, USA), and a solution of 100 U/ml penicillin and 100 *µ*g/ml streptomycin (GibcoBRL, Grand Island, NY, USA), and grown at 37°C in a 5% CO_2_ incubator at physiological pH. Cells were seeded on 25 cm2 cell culture T flasks (Sarstedt, Numbrecht, Germany) and sub-cultured by trypsin-EDTA (GibcoBRL, Grand Island, NY, USA) treatment. During the experiments reported below the medium was not refreshed.

#### Growth rate/Cell proliferation assay

Proliferation assays were conducted in 25 cm2 T flasks, starting with a cell density of 8.2×105 and 5.2×105 cells/ml for Huh7 and PLC, respectively. At the time points indicated, media were collected, cells were washed and trypsinized with a 0.25% (W/V) solution of trypsin (GibcoBRL, Grand Island, NY, USA). The total number of cells in the consequent supernatant was determined by hemocytometer counting (viable plus non-viable). Mean growth rate was determined by counting six non-overlapping sets of sixteen corner squares selected at random, and these four times at each time point.

#### Metabolic assays

The concentrations of glucose, glutamine and lactate in samples from the cells’ supernatant were determined by High Performance Liquid Chromatography (HPLC) based on calibration curves made with standard solutions. The samples were taken from supernatant at the end of experiment, filtered using 0.22*µ*m syringe filters (BGB Analytik Vertrieb GmbH, Rheinfelden, Germany) and stored until measurement in -80°C. HPLC was performed using the HPLC-DAD RID LC-20AT Prominence (Shimadzu, Columbia, USA) machine with a UV Diode Array Detector SPD-M30A NexeraX2 or/and a Refractive Index Detector RID 20A and an analytical ion-exclusion Rezex ROA-Organic Acid H+(8%) column (250×4.6 mm) with guard column (Phenomenex, Torrance, USA) (5 mM H2SO4 in MilliQ water (18.2 MΩ), isocratic, 0.15 ml/min. flow rate). Injection volume was 15 *µ*l (Autosampler: SIL-20AC, Prominence, Shimadzu), column oven temperature was 55°C (Column oven: CTO-20A, Prominence, Shimadzu) and the pressure was 29 bar.

## Author Contributions

conceptualization, E.W-T.; methodology, E.W-T.,T.D.M and H.V.W; software, T.D.M; experimental work, E.W-T and D.P.; validation, T.D.M. and E.W-T.; formal analysis, E.W-T.,T.D.M and H.V.W; investigation, E.W-T.,T.D.M and H.V.W; resources, T.D.M and E.W-T.; writing–original draft preparation, E.W-T.,T.D.M and H.V.W; writing–review and editing, E.W-T.,T.D.M and H.V.W; visualization, E.W-T. and T.D.M; supervision, H.V.W.; project administration, E.W.-T.

## Acknowledgments

This work is co-supported by the research Priority Area of the University of Amsterdam. We gratefully acknowledge Eugenie Troia and Hugo Pineda Hernández for their kind help with HPLC and Stefania Astrologo for pointing out the existing datasets. We thank Chiara Damiani for discussing possible extensions of the MaREA methodology. We are thankful to Jakub M. Tomczak for helping out at the final stage of preparing the manuscript.

