## Supplementary for "Simultaneous integration of gene expression and nutrient availability for studying metabolism of hepatocellular carcinoma"

### Supporting Information
### for

Simultaneous integration of gene expression and nutrient availability for studying metabolism of hepatocellular carcinoma

Ewelina Węglarz-Tomczak[1*], Thierry D.G.A. Mondeel[1], Diewertje G.E. Piebes[1], Hans V. Westerhoff[1,2,3]

[1]Swammerdam Institute for Life Sciences, Faculty of Science, University of Amsterdam, Amsterdam, The Netherlands
[2]Molecular Cell Physiology, Amsterdam Institute for Molecules, Medicines and Systems, Faculty of Science, Vrije Universiteit Amsterdam, Amsterdam, The Netherlands
[3]Manchester Centre for Integrative Systems Biology, School for Chemical Engineering and Analytical Sciences, University of Manchester, Manchester, UK

*Corresponding author


#### S1. Supplementary figures

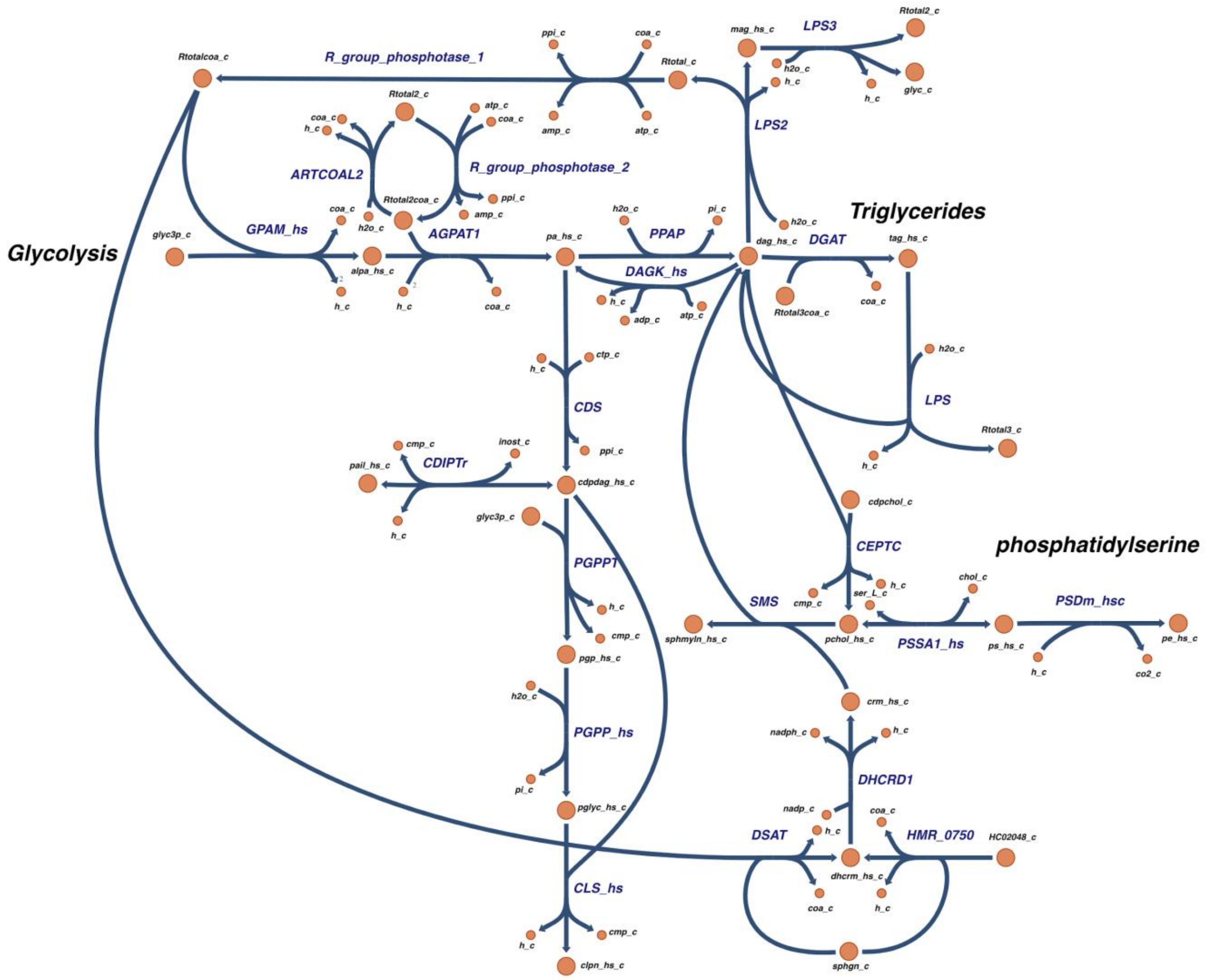

**Figure S1.** Network diagram of the subnetwork of the published version of Recon3.01 related to lipid synthesis from glycerol-3-phosphate (referred to as *glyc3p_c*) drawn using Escher [1]. See Figure S2 for the updated network diagram.

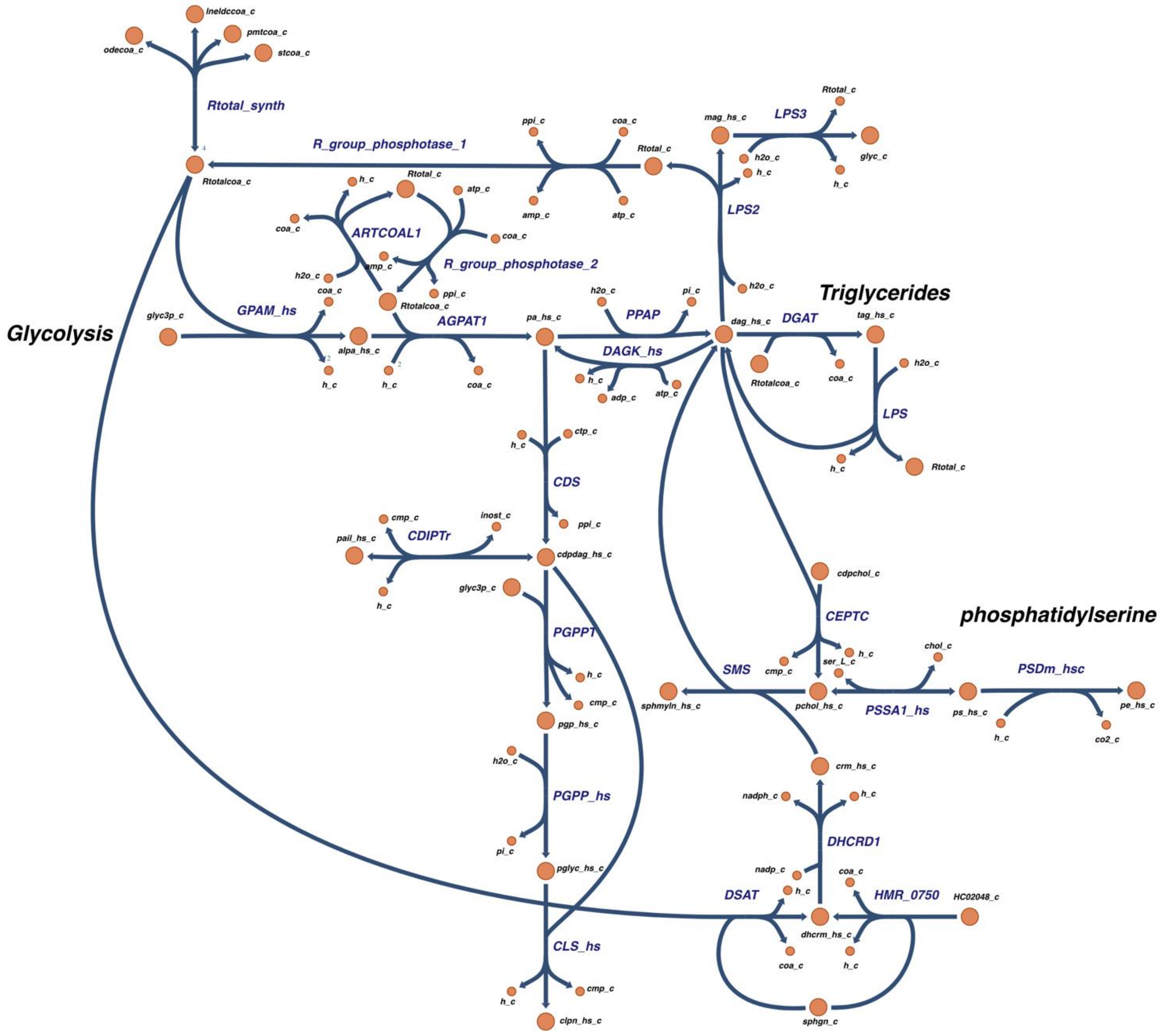

**Figure S2.** Network diagram of the updated subnetwork in Recon3.01 related to lipid synthesis from glycerol-3-phosphate (referred to as *glyc3p_c*) drawn using Escher [1]. We enhanced the pathway by adding a synthesis reaction for Rtotal and by equating the various forms of Rtotal by adding reversible reactions between them (see main text).

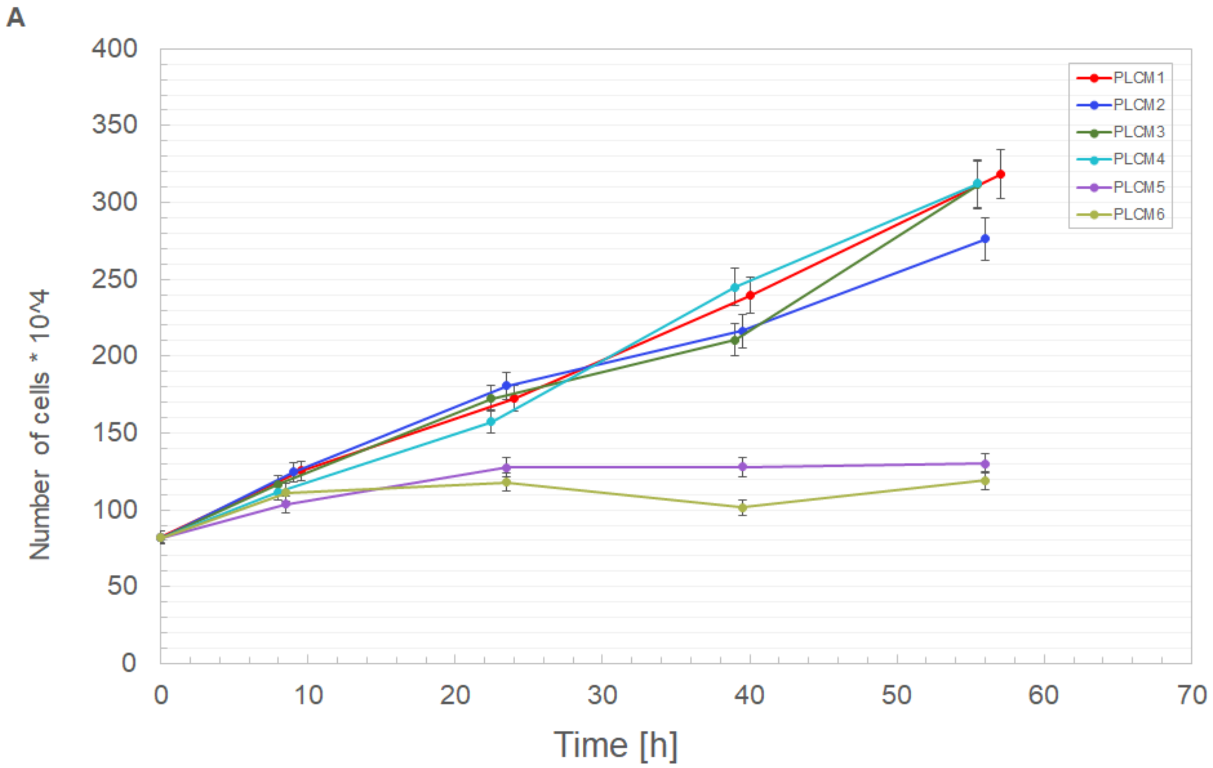
A
Number of cells * 10^4
Time [h]
PLCM1
PLCM2
PLCM3
PLCM4
PLCM5
PLCM6

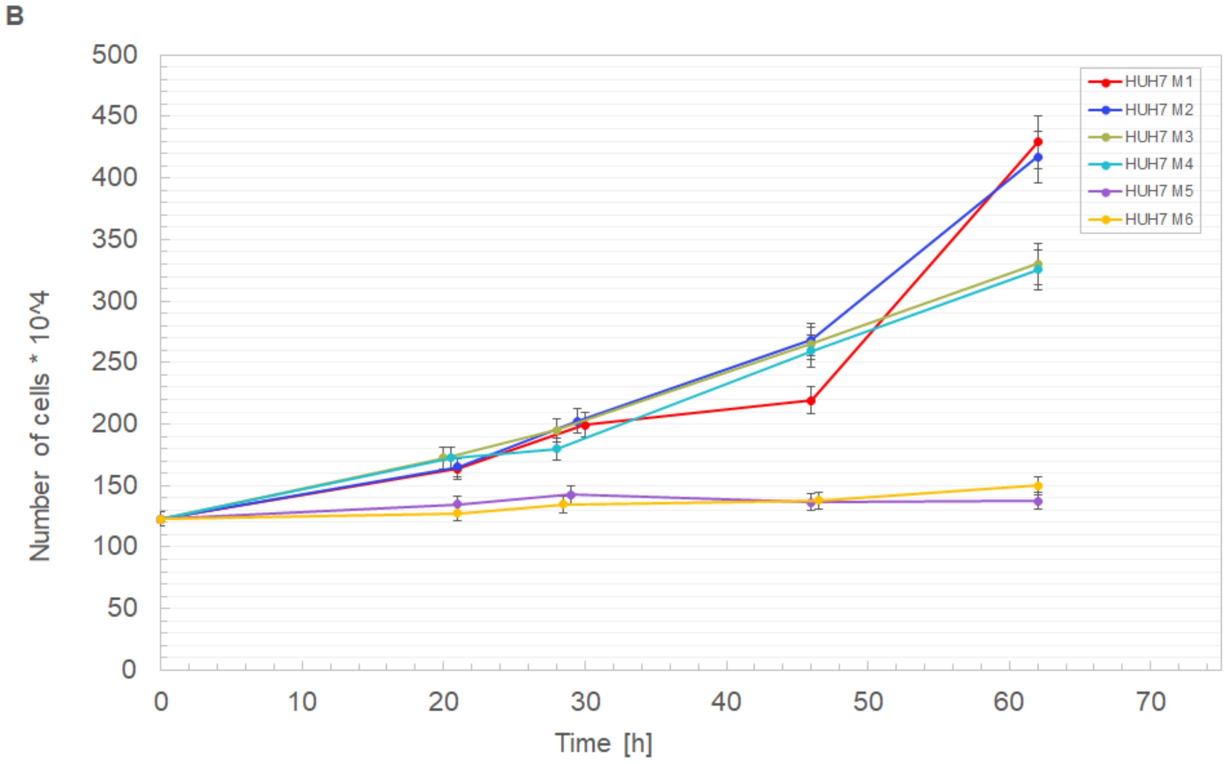
B
Number of cells * 10^4
Time [h]
HUH7 M1
HUH7 M2
HUH7 M3
HUH7 M4
HUH7 M5
HUH7 M6

C

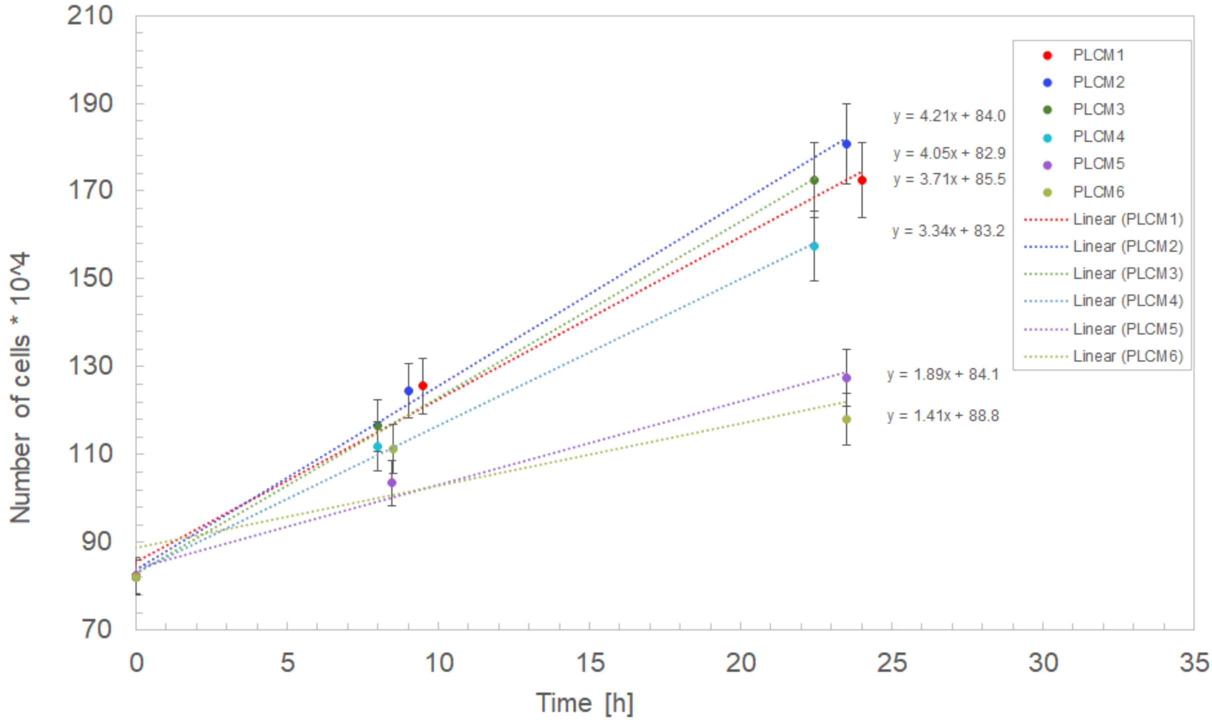

D

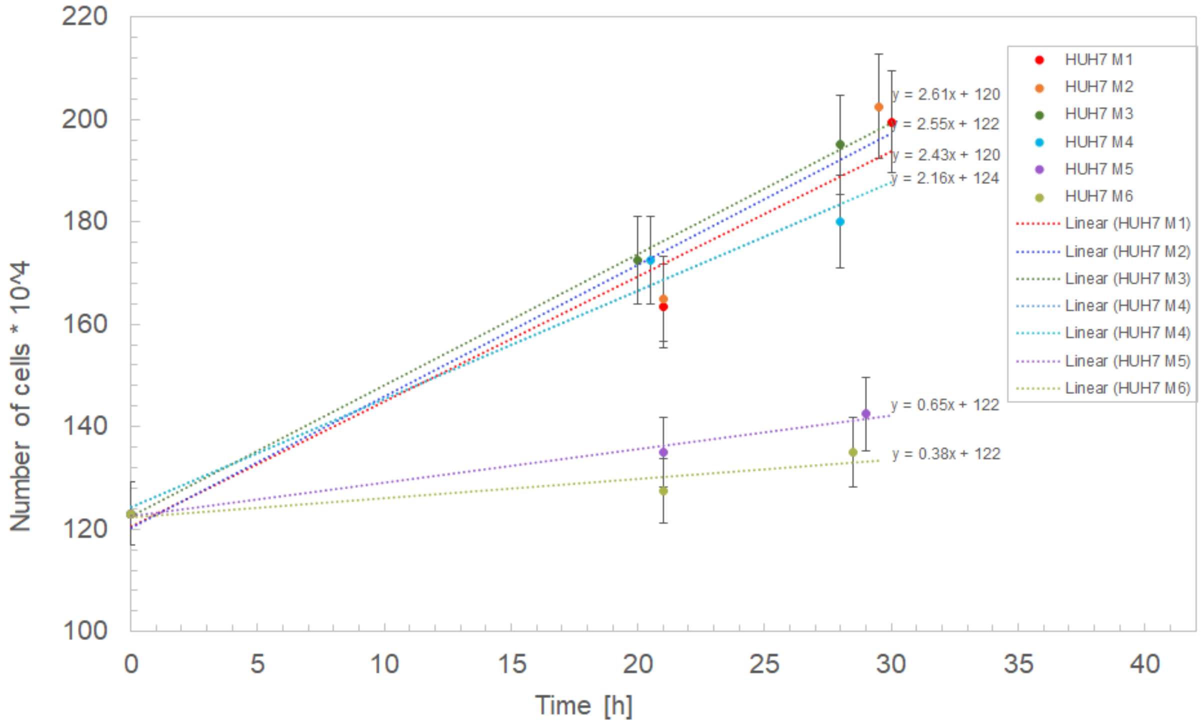

**Figure S3.** The growth rate of PLC (**A** and **C**) and Huh7 (**B** and **D**) cells in different media for the entire experiment (**A** and **B**) and the first 24 hours (**C** and **D**) that were used to calculate the linear relationship. The Y-axis represents the number of cells, the X-axis represents the time expressed in hours.

#### S2. Supplementary tables

**Table S1.** List of Entrez identifiers for genes present in Recon3D that we were not able to identify in the RNA-seq dataset by Ma et al. after converting the Entrez ID to the respective gene symbol and possible aliases. Red cells concern Entrez identifiers that do not exist, whereas orange cells concern Entrez identifiers that have been withdrawn from NCBI. The 16 genes in white or orange cells were assumed to be expressed at excess levels.

| Entrez ID | Symbol | Alias | Name |
|---|---|---|---|
| 201288.1 | NOS2P2 | NOS2B | nitric oxide synthase 2 pseudogene 2 |
| 645740.1 | NOS2P1 | NOS2C | nitric oxide synthase 2 pseudogene 1 |
| 728441.1 | GGT2 | GGT, GGT 2 | gamma-glutamyltransferase 2 |
| 102724560.1 | CBSL | CBS | cystathionine-beta-synthase like |
| 65263.1 | PYCR3 | PYCRL | pyrroline-5-carboxylate reductase 3 |
| 8781.1 | PSPHP1 | CO9, PSPHL | phosphoserine phosphatase pseudogene 1 |
| 100507855.1 | | | |
| 644378.1 | GCNT2P 1 | GCNT2P, GCNT6 | GCNT2 pseudogene 1 |
| 284004.1 | HEXD | HEXDC | hexosaminidase D |
| 8041.1 | | | |
| 100288072.1 | SDR42E 2 | | short chain dehydrogenase/reductase family 42E, member 2 |
| 340811.1 | AKR1C8 P | AKR1CL1 | aldo-keto reductase family 1 member C8, pseudogene |
| 0 | | | |
| 4514.1 | COX3 | COIII, MTCO3 | cytochrome c oxidase III |

| | | | |
|---|---|---|---|
| 4513.1 | COX2 | COII, MTCO2 | cytochrome c oxidase subunit II |
| 4512.1 | COX1 | COI, MTCO1 | cytochrome c oxidase subunit I |

**Table S2.** Metabolic components of the biomass reaction in Recon3D listed along with the stoichiometry at which they were taken to contribute to the biomass pseudo reaction. The stoichiometric coefficient is negative if the compound was taken to be consumed in the biomass synthesis reaction and positive if it was produced therein. It has been obtained [2] by determining the mass-fractional contributions of each biomass precursor to the dry weight of the biomass (i.e. in terms of milligram precursor per gram of dry weight (DW)). This fraction is then divided by the molecular weight of that precursor to obtain the above coefficient, which then is in unit mmol precursor per gram of biomass dry weight (g DW). The biomass reaction equation is then formulated as the sum of each metabolite precursor multiplied by its above coefficient. The biomass reaction rate is the specific growth rate and has units (g DW / g DW)/h. The specific consumption rates of the metabolites are obtained by multiplying their above coefficients by the specific growth rate and thereby are in units mmol/(g DW)/h [3]. Multiplying these specific consumption rates by the biomass concentration in g DW/dm$^3$ of the culture vessel, one obtains the consumption flux in terms of mM/h.

| **Metabolite** | **Coefficient** |
|---|---|
| Adenosine Diphosphate | 20.65082 |
| L-Alanine | -0.50563 |
| L-Arginine | -0.35926 |
| L-Asparagine | -0.27943 |
| L-Aspartate | -0.35261 |
| Adenosine Triphosphate | -20.7045 |
| Cholesterol | -0.0204 |

| | |
|---|---|
| Cardiolipin | -0.01166 |
| Cytidine-5'-Triphosphate | -0.03904 |
| L-Cysteine | -0.04657 |
| Deoxyadenosine Triphosphate | -0.01318 |
| Deoxycytidine-5'-Triphosphate | -0.00944 |
| Deoxyguanosine-5'-Triphosphate | -0.0099 |
| Deoxythymidine-5'-Triphosphate | -0.01309 |
| D-Glucose 6-Phosphate | -0.27519 |
| L-Glutamine | -0.326 |
| L-Glutamate | -0.38587 |
| Glycine | -0.53889 |
| Guanosine-5'-Triphosphate | -0.03612 |
| Water | -20.65082 |
| Proton | 20.65082 |
| L-Histidine | -0.12641 |
| L-Isoleucine | -0.28608 |
| L-Leucine | -0.54554 |
| L-Lysine | -0.59211 |
| L-Methionine | -0.15302 |
| 1-Phosphatidyl-1D-Myo-Inositol | -0.02332 |

| | |
|---|---|
| Phosphatidylcholine | -0.15446 |
| Phosphatidylethanolamine | -0.05537 |
| Phosphatidylglycerol | -0.00291 |
| L-Phenylalanine | -0.25947 |
| Orthophosphate | 20.65082 |
| L-Proline | -0.41248 |
| Phosphatidylserine | -0.00583 |
| L-Serine | -0.39253 |
| Sphingomyelin | -0.01749 |
| L-Threonine | -0.31269 |
| L-Tryptophan | -0.01331 |
| L-Tyrosine | -0.15967 |
| Uridine-5'-TrIphosphate | -0.05345 |
| L-Valine | -0.35261 |

**Table S3.** List of the oxidative phosphorylation reactions considered in Figure 5 by their reaction ID in Recon3D and with their reaction as specified in Recon3D.

| **ID** | **Name** | **Reaction** |
|---|---|---|
| ATPS4mi | ATP synthase | adp_m + 4.0 h_i + pi_m --> atp_m + h2o_m + 3.0 h_m |
| NADH2_u10mi | NADH dehydrogenase | 5.0 h_m + nadh_m + q10_m --> 4.0 h_i + nad_m + q10h2_m |

| | | |
|---|---|---|
| CYOR_u10mi | ubiquinol-cytochrome c reductase | 2.0 ficytC_m + 2.0 h_m + q10h2_m --> 2.0 focytC_m + 4.0 h_i + q10_m |
| CYOOm2i | cytochrome c oxidase | 4.0 focytC_m + 8.0 h_m + o2_m --> 4.0 ficytC_m + 2.0 h2o_m + 4.0 h_i |
| CYOOm3i | cytochrome c oxidase | 4.0 focytC_m + 7.92 h_m + o2_m --> 4.0 ficytC_m + 1.96 h2o_m + 4.0 h_i + 0.02 o2s_m |
| PDHm | pyruvate dehydrogenase | coa_m + nad_m + pyr_m --> accoa_m + co2_m + nadh_m |

**Table S4.** Concentrations of metabolites in the DMEM media in mM as listed in the manufacturer's formulation. Our media M1-M6 only differ in the glucose and glutamine concentrations (colored in green) and were exempt of ammonia. All concentrations were effected in-silico as maximal uptake rates which shape and constrain the flux cone of the solutions in FBA (see text). For carbon dioxide and oxygen, a not limiting uptake bound of 1000 was taken. Oxygen was considered non-limiting because the concentration in the medium was in the order of 0.2-0.4 mM (corresponding to air saturated saline) [4] whereas the $K_m$ of cytochrome oxidase for oxygen is some 0.01 mM [5]. Carbon dioxide was considered non-limiting due to its continuous replenishment in the medium.

| **Metabolite** | **Uptake bound** | **Metabolite** | **Uptake bound** | **Metabolite** | **Uptake bound** | **Metabolite** | **Uptake bound** |
|---|---|---|---|---|---|---|---|
| Arginine | 0.40 | Bicarbonate | 44 | Nicotinamide | 0.033 | Thiamin | 0.011 |
| Choline | 0.028 | Histidine | 0.2 | Valine | 0.80 | Threonine | 0.80 |
| Cysteine | 0.20 | Isoleucine | 0.80 | Phenylalanine | 0.4 | Tryptophan | 0.078 |
| Iron (Fe3+) | 0.00024 | Myo-Inositol | 0.04 | Phosphate | 0.9 | Tyrosine | 0.40 |
| Folate | 0.0091 | Potassium | 5.3 | (R)-Pantothenate | 0.0083 | Carbon dioxide | 1000 |
| Glycine | 0.40 | Methionine | 0.20 | Serine | 0.40 | Oxygen | 1000 |
| Water | [54220, | Sodium | 155 | Sulfate | 0.81 | | |

| | | | | | |
|---|---|---|---|---|---|
| | 54250, 54100, 54440, 54470, 54500] | | | | |
| Glucose | [0,5.6,25] | Leucine | 0.80 | Pyridoxine | 0.019 |
| Glutamine | [0, 4.0] | Lysine | 0.80 | Riboflavin | 0.0011 |
